# Targeting the expansion of myeloid-derived suppressor cells in liver cirrhosis

**DOI:** 10.1101/2024.03.29.587228

**Authors:** Emilio Flint, Caner Ercan, Eoin Mitchell, Oltin T Pop, Anne Geng, Paul OG Jorzik, Lucia Possamai, Robert G Brenig, Sarah Morel, Pablo Sieber, Arjuna Singanayagam, Matthias S Matter, David Semela, Markus H Heim, Philippe Demougin, Julien Roux, Luigi M Terracciano, Evangelos Triantafyllou, Christine Bernsmeier

## Abstract

**Background and aims:** Previously, we identified immune-suppressive circulating monocytic myeloid-derived suppressor cells (M-MDSC) in patients with cirrhosis and liver failure, which increased with disease severity and were associated with infections and mortality. Impaired immune responses and M-MDSC expansion were reversed by *ex vivo* polyinosinic:polycytidylic acid (poly(I:C)) treatment. Here, we aimed to investigate hepatic MDSC subsets in liver biopsies of cirrhotic patients and identify MDSC subsets in murine models to assess the safety and efficacy of poly(I:C) *in vivo*.

**Methods:** 22 cirrhotic patients and 4 controls were clinically characterised. MDSC were identified in liver biopsies (immunofluorescence) and in the circulation (flow cytometry). M-MDSC phenotype and function following poly(I:C) stimulation were assessed *ex vivo*. Carbon tetrachloride-based murine models of liver fibrosis were used. Poly(I:C) was administered therapeutically. MDSC biology was investigated with flow cytometry, immunofluorescence and T-cell proliferation assay. Hepatic histopathology, transcriptomics (BulkRNAseq) and serum markers were assessed.

**Results:** Besides circulating M-MDSC, hepatic CD14^+^CD84^+^M-MDSC and CD15^+^CD84^+^ polymorphonuclear-MDSC expanded in cirrhotic patients and indicated disease severity, infections and poor survival. Poly(I:C) treatment reversed phenotype and function of circulating M-MDSC *ex vivo*. Circulating and hepatic MDSC expanded in our murine models of liver fibrosis and suppressed T-cell proliferation. Lipopolysaccharide and *E.coli* challenge exacerbated hepatic MDSC and fibrosis compared to CCl_4_ controls. Poly(I:C) therapy reduced MDSC expansion in fibrotic mice with bacterial infection and CCl_4_-induced fibrosis.

**Conclusion:** Hepatic MDSC expanded in cirrhotic patients and were linked with disease severity and poor prognosis. Poly(I:C) reversed frequency and function of M-MDSC *ex vivo*. Poly(I:C) therapy reversed MDSC expansion and fibrosis in a murine model of liver fibrosis with infection. Thus, we highlighted poly(I:C) as a potential immunotherapy for the treatment of immuneparesis in cirrhosis.

## Introduction

Patients with cirrhosis are at a high risk of developing infections[1], occurring five to six times more frequently than in hospitalized patients overall[2]. Infections drive hepatic decompensation and are associated with increased rates of hospitalisation, extrahepatic organ failure and death, lowering the 1-year transplant-free survival to less than 60%[3–5].The pathophysiology underlying increased infection susceptibility in patients with cirrhosis is complex, and involves a wide range of cellular and soluble factors across multiple compartments of the body, which have been collectively termed immuneparesis[6, 7].

MDSC are pathological subsets of monocytes and neutrophils that strongly suppress immune responses, including those mediated by T cells, B cells and natural killer cells[8]. In both humans and mice, two subsets of MDSC can be distinguished based on their myeloid lineage, i.e. monocytic (M-MDSC) and polymorphonuclear (PMN-MDSC) MDSC[8]. These cells arise in a variety of pathological conditions, including the tumour microenvironment of cancers[9], chronic inflammatory- [10], autoimmune disorders[11], chronic viral infections[12] (including hepatitis B and C[13, 14]), sepsis[15] as well as cirrhosis and acute decompensation (AD) or acute-on-chronic liver failure (ACLF)[16].

Previously, we detailed the expansion of circulating M-MDSC during progression of cirrhosis, which peaked in liver failure, representing 55% of circulating CD14^+^ cells, and was associated with secondary infections, disease severity and poor prognosis[16]. While M-MDSC expansion has been detailed in the circulation of patients with cirrhosis and AD/ACLF[16], the presence and distribution of MDSC subsets in the liver of cirrhotic patients remains unknown. In this study, we sought to investigate the frequency of hepatic MDSC subsets in liver biopsies from patients with cirrhosis in relation to prognostic clinical parameters. In parallel, we investigated the presence and distribution of MDSC subsets in carbon tetrachloride (CCl_4_) induced murine models of liver fibrosis.

Another aim of our study was to assess both immunotherapeutic effects and liver-related safety parameters following poly(I:C) administration in a CCl_4_ model. We previously showed that impaired anti-microbial innate immune responses and expansion of M-MDSC were reversed *ex vivo* by TLR-3 agonism with polyinosinic:polycytidylic acid (poly(I:C))[16]. Other studies also showed that poly(I:C) promoted phagocytic activity by macrophages[17], enhanced infection clearance and survival in a murine pneumonia model[18], reduced MDSC frequency in the tumour microenvironment of liver metastases[19–23] and attenuated hepatic fibrosis in multiple murine models of liver fibrosis[24–26]. Poly(I:C) may thus represent an interesting candidate for potential future immunotherapy in cirrhosis. However, the effects of poly(I:C) immunotherapy on MDSC expansion, innate immune responses and liver-related safety parameters in preclinical models of cirrhosis remain unknown.

## Results

### Hepatic MDSC subsets expanded with disease progression in patients with cirrhosis

In order to explore whether MDSC expand in the livers of patients with cirrhosis, we investigated tissue from liver biopsies of 22 cirrhotic patients, categorized into Child-Pugh classes A (n = 8), B (n = 7), and C (n = 7), as well as a control group of 4 individuals without liver disease. The clinical features of these patients are summarised in Table 1. Patient characterization was conducted based on aetiology, disease severity scores as well as a broad range of clinical parameters (Table 1). Within a one-year time frame, transplant-free survival was 72.7% (n = 16), orthotopic liver transplantation was conducted in 13.6% of patients (n = 3) and the mortality rate was 13.6% (n = 3) distributed among ACLF 33% (n = 1), sepsis 33% (n = 1) and pneumonia 33% (n = 1). Infections occurred in 27.3% of patients (n = 6), including spontaneous bacterial peritonitis 50% (n = 2), urinary tract infections 50% (n = 2), hospital-acquired pneumonia 25% (n = 1) and sepsis of unidentified origin 25% (n = 1).

**Table 1.** Clinical characteristics of patients with cirrhosis at different Child Pugh stages compared to individuals without liver disease. NA, not applicable; ALD, alcoholic liver disease; HCV, hepatitis C virus; NAFLD, Non-alcoholic fatty liver diseas; PBC, primary biliary cirrhosis; MELD, model for end-stage liver disease; BMI, body mass index; AST, aspartate aminotransferase; ALT, alanine aminotransferase; GGT, gamma-glutamyl transferase; AP, alkaline phosphatase; INR, international normalized ratio; CRP, C-reactive protein. Data indicated as median with (minimum and maximum) or [percentage]. *P < 0.05 indicates Child A vs Child B; #P < 0.05, ##P < 0.01 indicate Child-Pugh A vs Child-Pugh C; †P < 0.05 indicates Child-Pugh B vs Child-Pugh C, comparisons by Mann-Whitney U Tests.

| Variables<br>median (range) | Individuals without liver<br>disease (n=4) | Child-Pugh A (n=8) |  | Child-Pugh B (n=7) |  | Child-Pugh C (n=7) |  |
| --- | --- | --- | --- | --- | --- | --- | --- |
| Age (years) | 55 (47–69) | 56 (44–75) |  | 74 (56–79) |  | 58 (34–77) |  |
| Sex (m:f) | - | 3:5 |  | 5:2 |  | 5:2 |  |
| Underlying liver<br>disease(s) [%] | NA | ALD | 6 [75] | ALD | 2 [26] | ALD | 3 [43] |
|  |  | HCV | 2 [25] | PBC | 2 [26] | HCV | 1 [14] |
|  |  |  |  | HCV | 1 [14] | NAFLD | 1 [14] |
|  |  |  |  | NAFLD | 1 [14] | Wilson’s disease | 1 [14] |
|  |  |  |  | Unknown | 1 [14] | Hemochromatosis | 1 [14] |
| Child-Pugh | NA | 6 (5–7) |  | 8 (7–9) |  | 10 (10–11) |  |
| MELD | NA | 8 (6–10)<sup>##</sup> |  | 10 (6–21)<sup>†</sup> |  | 16 (15–29) |  |
| BMI | - | 27.3 (19.7–31.2) |  | 24 (17.1–30.1) |  | 28.2 (27.4–31.9) |  |
| Na<sup>+</sup> (mmol/l) | 140 (138–142) | 140 (137–141) |  | 140 (136–144) |  | 136 (120–144) |  |
| K<sup>+</sup> (mmol/l) | 3.95 (3.40–4.20) | 3.85 (3.3–4.4) |  | 4.1 (3–5.1) |  | 3.9 (3.3–4.4) |  |
| Creatinin<br>(μmol/l) | 83.5 (73.0–93.0) | 58.5 (50–63)<sup>*</sup> |  | 93 (65–171) |  | 78 (47–165) |  |
| Urea (mmol/l) | 6.75 (4.8–7.6) | 4.9 (2.4–6.6) |  | 7.6 (5.6–14.7) |  | 4.1 (3.1–7.6) |  |
| Bilirubin<br>(μmol/l) | 9.5 (6–12) | 14.5 (3–49)<sup>##</sup> |  | 17 (8–77) |  | 64 (21–147) |  |
| AST (U/l) | 23 (22–27) | 38 (27–57)<sup>#</sup> |  | 36 (16–111) |  | 65 (39–168) |  |
| ALT (U/l) | 30.5 (15.0–35.0) | 30 (11–43) |  | 20 (7–51) |  | 52 (11–114) |  |
| GGT (U/l) | 119 (20–133) | 141.5 (41–530) |  | 103 (28–199) |  | 61 (26–1013) |  |
| AP (U/l) | 56 (33–93) | 78.5 (56–128) |  | 141 (74–257) |  | 115 (49–483) |  |
| INR | - | 1.0 (1.0–1.30)<sup>#</sup> |  | 1.1 (0.9–2.3) |  | 1.6 (1.4–2.3) |  |
| Albumin (g/l) | 39 (38–40) | 28.5 (20–39.0) |  | 30.6 (19–30.9) |  | 25 (19.2–30.1) |  |
| Hemoglobin<br>(g/l) | - | 140.5 (81–171) |  | 104 (84–124) |  | 98 (7.2–116) |  |
| Hematocrit (%) | - | 0.42 (0.3–0.49)<sup>#</sup> |  | 0.31 (0.26–0.38) |  | 0.27 (0.24–0.35) |  |
| Leucocytes (G/l) | - | 7.3 (3.2–11.6) |  | 3.0 (2.4–8.1) |  | 5.3 (3.8–70) |  |
| Neutrophils<br>(G/l) | - | 56.6 (40.4–80.6) |  | 67.6 (53.1–71.5) |  | 59 (55.3–83.2) |  |
| Eosinophils (G/l) | - | 2.9 (1.3–4.7) |  | 5.5 (1.3–10.4) |  | 1.3 (0.1–8.6) |  |
| Basophils (G/l) | - | 0.6 (0.4–1.4) |  | 0.4 (0.3–0.5) |  | 0.4 (0.2–1.9) |  |
| Lymphocytes<br>(G/l) | - | 1.9 (1.3–2.2)<sup>*</sup> |  | 0.6 (0.4–1.9) |  | 2.00 (1.2–2.5) |  |
| Monocytes (G/l) | - | 7.3 (0.46–11.2) |  | 11.1 (9.7–12) |  | 10.7 (3.9–11.2) |  |
| Platelets (G/l) | - | 142 (98–341) |  | 87 (40–191) |  | 82 (8.5–299) |  |
| CRP (mg/l) | 1.35 (0.70–2.40) | 3.1 (0.4–27.5) |  | 8 (2–25) |  | 10 (3–67) |  |

We performed immunofluorescence stains on liver biopsies to investigate the presence and distribution of CD14^+^CD84^+^ M-MDSC and CD15^+^CD84^+^ PMN-MDSC[27] in the livers of patients with cirrhosis across Child Pugh classes A to C (fig. 1A and 1B). In parallel with the expansion of M-MDSC in the circulation during disease progression in cirrhosis[16, 28], we observed that both M-MDSC and PMN-MDSC expanded in the cirrhotic livers compared with controls without liver disease (1C and 1H). In addition, there was a marked expansion of M-MDSC in Child-Pugh stages B and C (fig. 1C), while PMN-MDSC expanded in Child-Pugh C class patients (fig. 1H). The expansion of both M-MDSC and PMN-MDSC were independently associated with the presence of ascites, development of infections and reduced 1-year transplant-free survival (fig. 1D and 1I). MDSC were also numerically higher in patients who were diagnosed with HCC within a one-year timeframe, although these trends were not statistically significant (fig. S1A). In addition, the presence of M-MDSC correlated positively with disease severity scores (Child-Pugh, MELD); INR, ALT and creatinine levels (fig. 1E). Similarly, the expansion of PMN-MDSC positively correlated with disease severity scores (Child-Pugh, MELD); INR; bilirubin and ALT levels (fig. 1J). When comparing the frequency of circulating and hepatic M-MDSC in our cirrhotic cohort (fig. 1F), the expansion of these cells in the circulation positively correlated with M-MDSC numbers in liver tissue from the same patients (fig. 1G).

**Figure 1.**
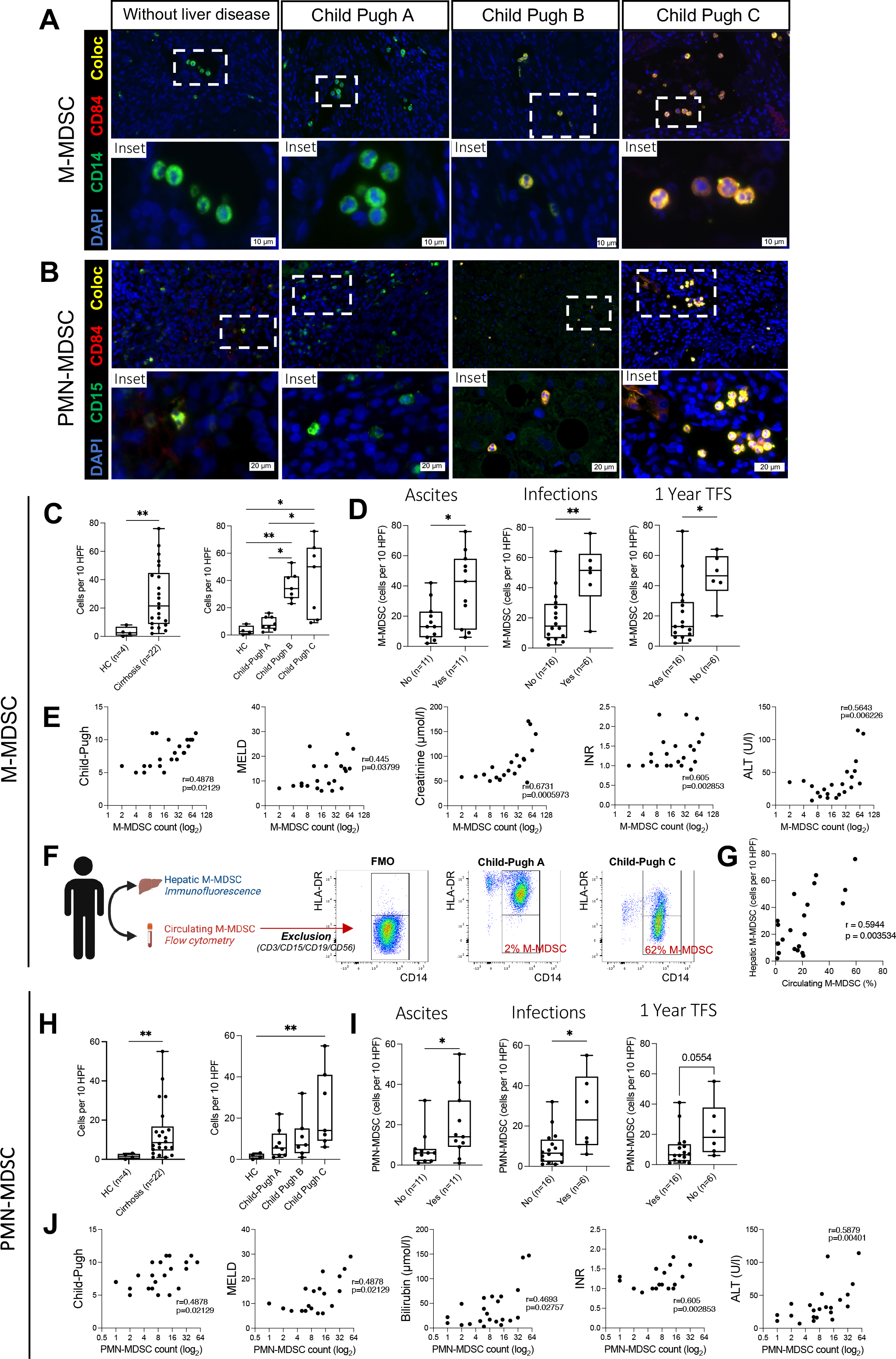
Hepatic MDSC subsets expand with disease progression in patients with cirrhosis. (A & B) Immunofluorescence micrographs of tissue sections from FFPE liver biopsies of healthy controls and patients with cirrhosis at stages Child-Pugh A to C. AF488=green depicts CD14 and CD15 expression (in panels A or B, respectively), AF647=red depicts CD84 expression, DAPI=blue indicates cell nuclei, scalebars indicate 10µm or 20µm (in panels A or B, respectively). **(C & H)** Quantification per 10 HPF magnification (400x) of CD14+CD84+ cells or CD15+CD84+ cells (in panels C or H, respectively) in healthy controls and patient with cirrhosis (n=22) and at stages Child-Pugh A to C. **(E & J)** Correlation plots with spearman r coefficients indicating correlations between M-MDSC counts or PMN-MDSC counts (in panels E or J, respectively) with disease severity scores Child-Pugh and MELD as well as laboratory parameters creatinine, ALT, bilirubin levels and INR. **(F)** Overview of experimental design and flow cytometry gating strategy for circulating M-MDSC in patients with cirrhosis **(G)** Correlation plot with spearman r coefficient indicating correlation between circulating M-MDSC (% of total monocytes) with M-MDSC counts from liver biopsies of the same patients. Box plots showing median with 10–90 percentile and all points min–max, *p < 0.05, **p < 0.01, statistical comparisons by Mann-Whitney tests (for n=2 groups) and Kruskal-Wallis test (for n>2 groups).

### Poly(I:C) stimulation *ex vivo* reduced the frequency of M-MDSC and restored the impaired innate immune function of monocytes

Flow cytometry analysis of either whole blood or isolated PBMCs from patients with cirrhosis identified CD14^+^CD15^-^HLA-DR^low/-^ M-MDSC in the circulation as previously described[16] (fig. 2A). Both M-MDSC and HLA-DR^int/hi^ monocytes expressed TLR3 (fig. 2B). In addition, M-MDSC expressed higher levels of CD84, a novel MDSC-specific marker[27, 29], compared to HLA-DR^int/hi^ monocytes (fig. 2C). Phagocytosis of both *E. coli* and *S. aureus* pHrodo bioparticles were impaired in M-MDSC compared to HLA-DR^int/hi^ monocytes (fig. 2D). In order to assess whether TLR3 agonism with poly(I:C) could reduce the frequency of M-MDSC and restore the impaired phagocytosis capacity of monocytes in patients with cirrhosis, we stimulated PBMCs isolated from cirrhotic patients with poly(I:C) (fig. 2E). TLR3 agonism markedly increased HLA-DR expression on monocytes and reduced the fraction of M-MDSC (fig. 2E and 2F). In addition, poly(I:C) stimulation reduced CD84 expression on monocytes overall (fig. 2G), while CD84 expression levels within M-MDSC and HLA-DR^int/hi^ subsets did not change following poly(I:C) stimulation (fig. S1B). Importantly, phagocytosis of *E. coli* and *S. aureus* particles by monocytes were increased upon stimulation with poly(I:C) (fig. 2H). Importantly, in other extrahepatic compartments such as peritoneal macrophages that were isolated from the ascites of patients with decompensated cirrhosis, we noted lower HLA-DR expression levels compared to monocyte-derived macrophages (fig. 2I). Finally, bone marrow tissue obtained from a cirrhotic patient prior to bone marrow transplantation displayed an expansion of CD14^+^CD84^+^ and CD15^+^CD84^+^ MDSC compared to control tissue (fig. 2J). Taken together, these data indicate that M-MDSC expand in the circulation of patients with cirrhosis and display impaired innate function compared to their healthy monocytic counterparts. TLR3 agonism via poly(I:C) reduced the frequency of MDSC and improved the innate immune function of monocytes from patients with cirrhosis.

**Figure 2.**
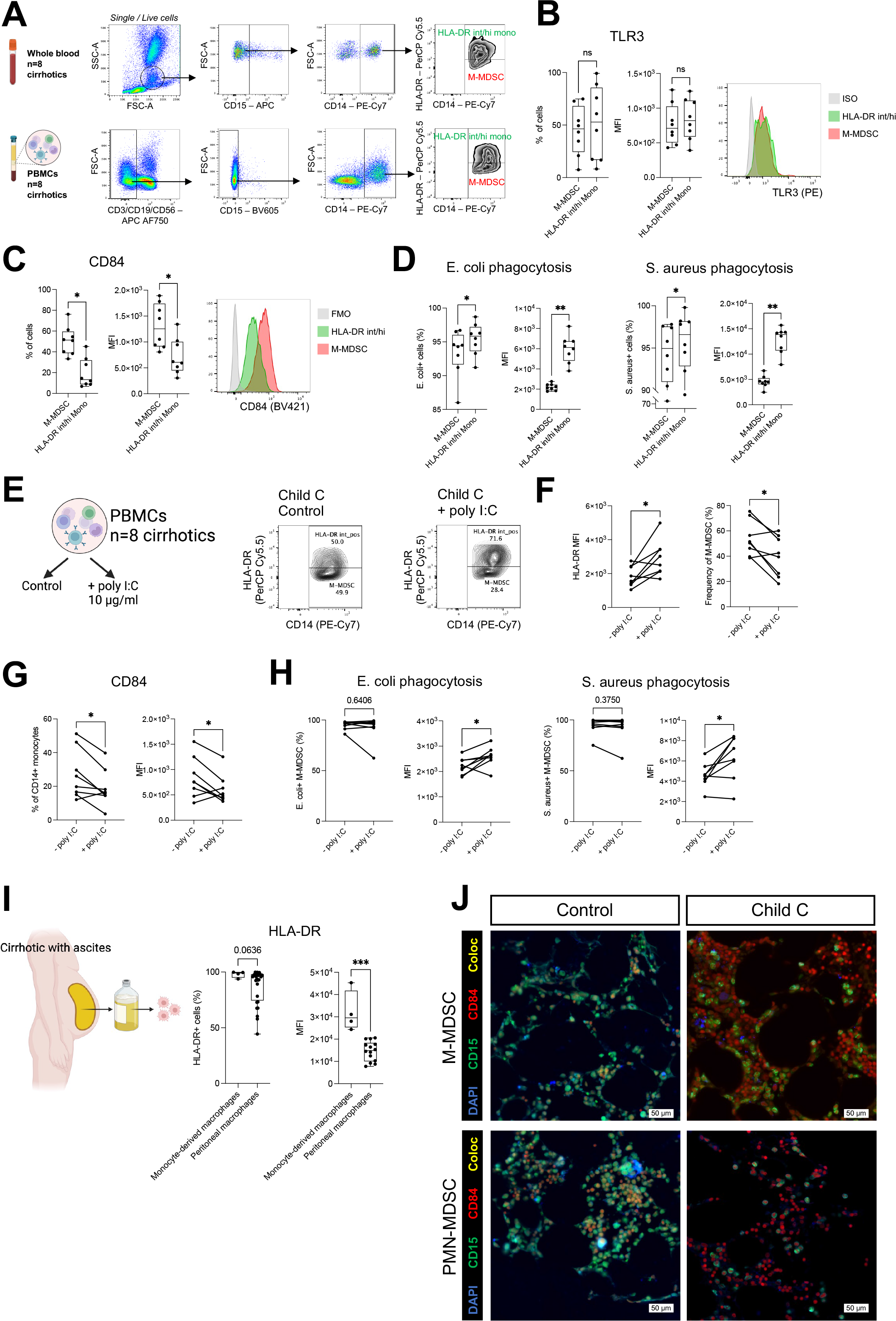
Poly(I:C) stimulation *ex vivo* reduced the frequency of M-MDSC and restored the impaired innate immune function of monocytes **(A)** Flow cytometry gating strategies for either M-MDSC or HLA-DR^int/hi^ monocytes from either whole blood or PBMCs from patients with cirrhosis. **(B and C)** Percentage and MFI values of TLR3 expression (panel B) and CD84 expression (panel C) on cell subsets identified in panel A. **(D)** Percentage and MFI values of PE-conjugated *E. coli* bioparticles or PE-conjugated *S. aureus* bioparticles in cell subsets **(E)** Design of *ex vivo* poly(I:C) stimulation of PBMCs and flow cytometry contour plots of CD14 / HLA-DR expression with or without poly(I:C) stimulation **(F)** HLA-DR MFI values and frequency of M-MDSC in 8 cirrhotic patients with or without poly(I:C) stimulation **(G)** Percentage and MFI values of CD84 expression on total monocytes from 8 cirrhotic patients with or without poly(I:C) stimulation. **(H)** Percentage and MFI values of PE-conjugated *E. coli* bioparticles or PE-conjugated *S. aureus* bioparticles in M-MDSC from 8 cirrhotic patients with or without poly(I:C) stimulation. **(I)** Schematic of ascitic tap from cirrhotic patient with ascites, as well as percentage and MFI values of HLA-DR expression in monocyte-derived macrophages (control) and peritoneal macrophages isolated from the ascites. **(J)** Immunofluorescence micrographs of tissue sections from FFPE bone marrow tissue obtained from a control subject and a cirrhotic patient prior to bone marrow transplantation. AF488=green depicts CD14/CD15 expression, AF647=red depicts CD84 expression, DAPI=blue indicates cell nuclei, scalebars=50µm. Box plots showing median with 10–90 percentile and all points min–max, *p < 0.05, **p < 0.01, ***p<0.001. Statistical comparisons by Mann-Whitney tests (unpaired data), Wilcoxon tests (paired data).

### MDSC subsets expand in a carbon tetrachloride-induced liver fibrosis model

In order to identify a suitable model to target the expansion of pathological MDSC subsets *in vivo*, we investigated the frequency and distribution of MDSC subsets in a murine model of fibrosis which involved intraperitoneal administration of CCl_4_, or vehicle control (olive oil), bi- weekly for a total of 6 weeks[30] (fig. 3A). Ishak Stage scoring and collagen proportionate area analysis of PicroSiriusRed (PSR) stains confirmed the establishment of liver fibrosis in our model (fig. 3B and 3C), and Ishak grading of H&E-stains indicated hepatic necroinflammation (fig. 3C). Moreover, these animals displayed reduced bodyweight gain compared to vehicle controls (fig. 3D). Accordingly, CCl_4_-treated mice had dramatically higher levels of blood ALT, indicating liver damage, while albumin blood levels were comparable to healthy controls, indicating no impairment in liver synthetic function (fig. 3E). While the weight of liver, kidney and spleen were unchanged in the CCl_4_ group, the liver to bodyweight ratio was increased in these animals (fig. 3F). Bulk RNA seq of murine liver tissue and pathway enrichment analysis revealed a variety of upregulated pathways following CCl_4_-treatment, including upregulation of extracellular matrix remodelling, collagen production as well as chemokines and chemokine receptors (fig. 3G).

**Figure 3.**
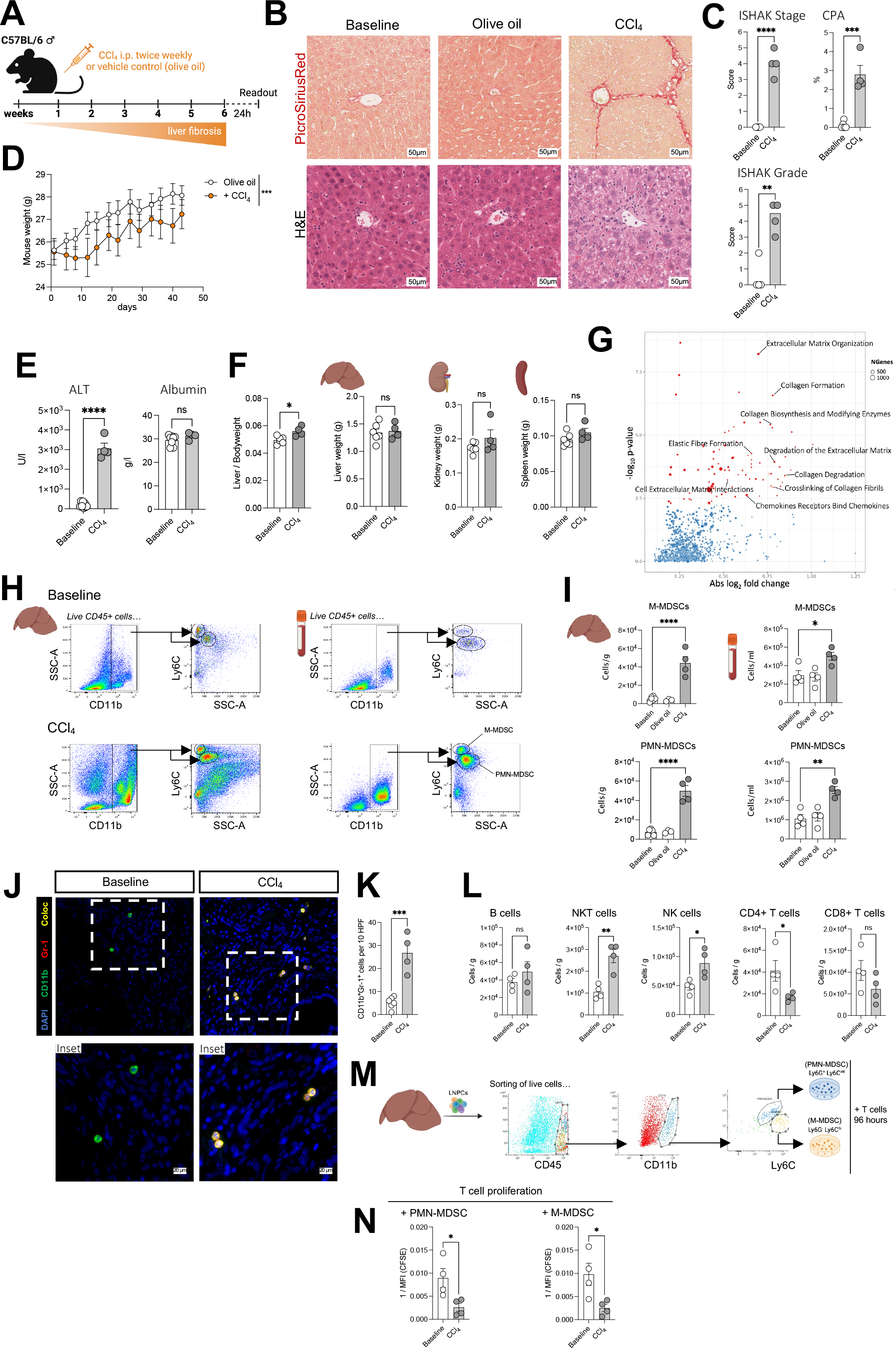
MDSC subsets expand in a carbon tetrachloride-induced liver fibrosis model. **(A)** Schedule of administration of CCl_4_ in C57BL/6 mice (duration 6 weeks). **(B)** Representative images of PicroSiriusRed (PSR) stains or hematoxylin & eosin (H&E) stains of liver sections from healthy age-matched controls, vehicle-treated controls (olive oil) or CCl_4_- treated mice, scalebar=100µm. **(C)** Ishak Stage and Ishak Grade scoring of stains from panel B, and CPA analysis of PSR stains. **(D)** Bodyweight curves over 6 weeks of CCl_4_ group (n=4) and vehicle-control group (n=4), statistical comparison by two-Way ANOVA mixed model. **(E)** Plasma levels of liver-related markers alanine aminotransferase and albumin **(F)** Liver-to-bodyweight ratio and total organ weight of liver, kidney and spleen from the CCl_4_ group and healthy controls. **(G)** Pathway enrichment analysis using gene sets from the Reactome database (significant at an FDR of 5% in red) comparing BulkRNAseq samples from liver tissue of CCl_4_-treated mice (n=4) vs healthy controls (n=6). **(H)** Flow cytometry gating strategies for M-MDSC and PMN-MDSC from single live CD45+ liver-non parenchymal cells (LNPC) or single live CD45+ blood cells, in healthy controls and CCl_4_-treated mice. **(I)** Quantification of M-MDSC or PMN-MDSC cells in liver and blood, normalized per gram of liver tissue or per millilitre of blood using flow cytometry counting beads. **(J)** Immunofluorescence micrographs of tissue sections from FFPE liver tissue from healthy controls and CCl_4_-treated mice. AF488=green depicts CD11b expression, AF647=red depicts Gr-1 expression, DAPI=blue depicts cell nuclei, scalebars=20µm. **(K)** Quantification per 10HPF magnification (400x) of CD11b+Gr-1+ cells (see panel I) **(L)** Quantification of lymphocyte subpopulations (gating strategy fig. S2D), normalised to gram of liver tissue. **(M)** Schematic depicting FACS of M-MDSC or PMN-MDSC from isolated LNPC from healthy controls and CCl_4_-treated mice followed by co-culturing with T cells **(N)** T cell proliferation following 96-hour co-culturing with cells (panel M) assessed via CFSE signal (population doublings). Barplots displaying all datapoints with mean ± SEM, statistical tests by unpaired t-tests (for n=2 groups) or one-way ordinary ANOVA with multiple comparisons (for n>2 groups), *p < 0.05, **p < 0.01, ***p < 0.001, ****p < 0.0001.

In order to assess the presence and distribution of MDSC subsets in this model, we isolated both liver non-parenchymal cells as well as blood cells and performed flow cytometry analyses. CCl_4_-treated mice displayed an expansion of CD45+ immune cells in the liver, indicating hepatic inflammation (fig. S2A). In both the liver and circulation, M-MDSC (CD45^+^CD11b^+^Ly6C^hi^SSC-A^int^) and PMN-MDSC (CD45^+^CD11b^+^Ly6C^low^SSC-A^hi^) expanded in CCl_4_-treated mice compared to untreated and vehicle controls (liver: M-MDSC 8.24-fold change p<0.0001, PMN-MDSC 6.61-fold change p<0.0001; blood: M-MDSC 1.70-fold change p=0.014, PMN-MDSC 2.41-fold change p=0.0012) (fig. 3H, 3I and S2B). In accordance with flow cytometry data, immunofluorescent stains revealed an expansion of CD11b^+^Gr-1^+^ tissue MDSC in the livers of CCl_4_-treated mice compared to untreated controls (5.177-fold change p=0.0003) (fig. 3J and 3K). In parallel to MDSC expansion, we noted a decrease in liver-resident Kupffer Cells (KC) (fig. S2C). Flow cytometry analysis of murine hepatic tissue identified lymphocyte subpopulations (fig. S2D). Interestingly, the number of CD4^+^ T cells was reduced in the livers of CCl_4_-treated mice (fig. 3L). To elucidate the immunosuppressive function of these cells, M-MDSC and PMN-MDSC were sorted from livers of either healthy controls or CCl_4_-treated mice using fluorescent-activated cell sorting and co-cultured with CFSE-labelled activated primary T cells. Accordingly, co-culturing with sorted M-MDSC or PMN-MDSC from livers of CCl_4_-treated mice reduced T cell proliferation (fig. 3M and 3N).

Taken together, these data indicate that immunosuppressive MDSC subsets expanded in the liver and circulation during CCl4-induced fibrosis, thereby supporting the use of this murine disease model to study the expansion and targetability of pathological MDSC *in vivo* during chronic liver injury.

### Hepatic MDSC expanded upon CCl_4_-induced fibrosis with either lipopolysaccharide stimulation or *E. coli* injection

In order to mimic the effects of recurrent bacterial challenge occurring in patients with cirrhosis, a group of CCl4-treated mice was administered 100 µg of lipopolysaccharide (LPS) intraperitoneally 24 hours prior to sacrifice (fig. 4A). Flow cytometry analysis revealed increased CD45+ immune infiltrates in the livers but not in the circulation of LPS-treated mice compared to CCl_4_ controls (fig. S3A). Interestingly, LPS challenge further expanded M-MDSC and PMN-MDSC in the liver, while numbers of MDSC subsets in the blood remained the same compared to CCl_4_ controls (liver: M-MDSC 1.78-fold change p=0.0031, PMN-MDSC 1.46-fold change p=0.041; blood: M-MDSC 1.17-fold change p=0.221, PMN-MDSC 1.05-fold change p=0.615) (fig. 4B, 4C and S3B). Immunofluorescence imaging of liver tissue revealed an expansion of CD11b^+^Gr-1^+^ cells in mice treated with CCl_4_ alone and CCl_4_ with LPS stimulation (fig. 4D and 4E). Remarkably, Ishak stage scoring and CPA analysis indicated that LPS administration exacerbated liver fibrosis and hepatic necroinflammation compared to CCl_4_ controls (fig. 4F and 4G).

**Figure 4.**
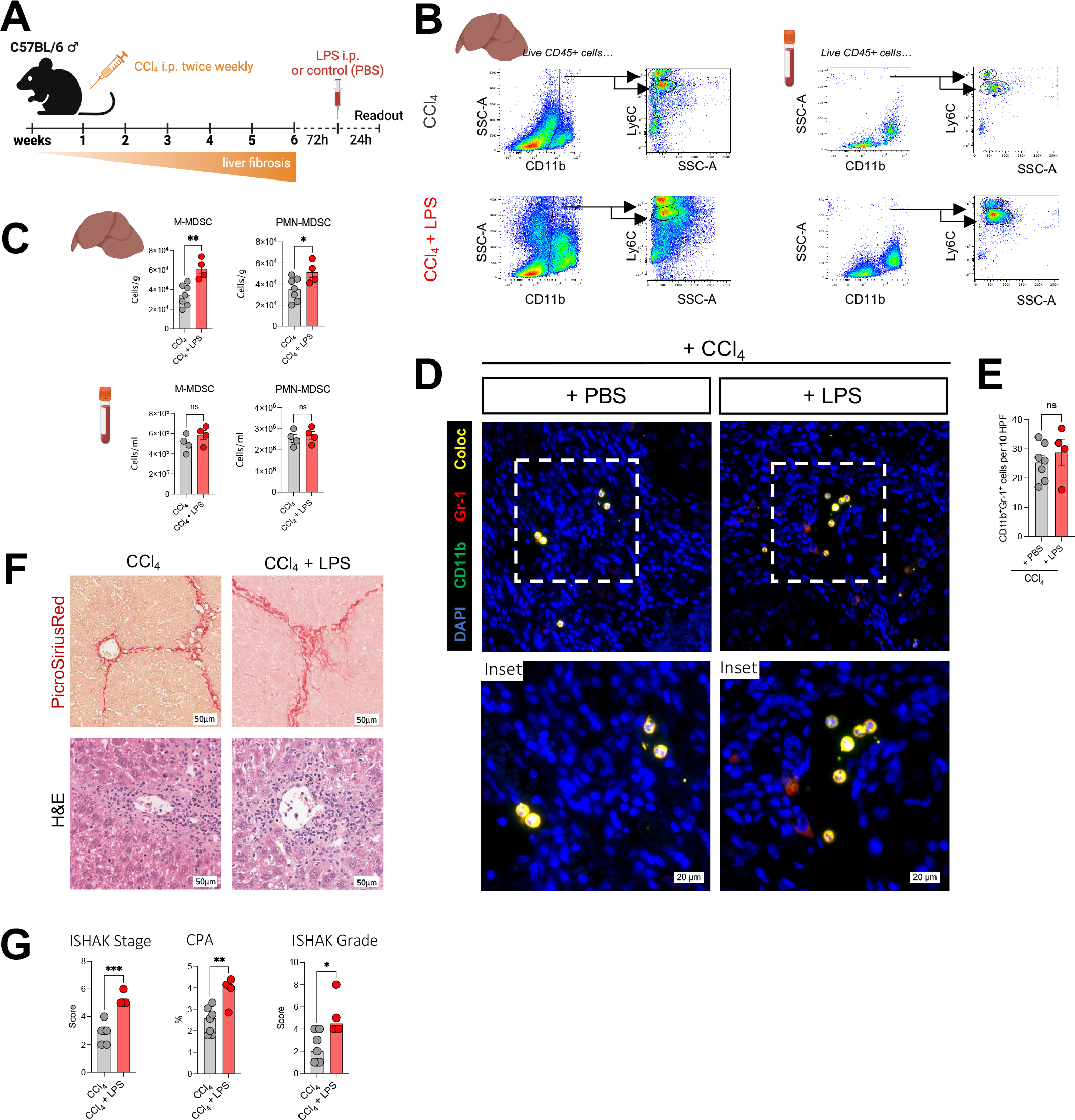
Lipopolysaccharide challenge further expands hepatic MDSC and exacerbates CCl_4_-induced fibrosis. (A) Schedule administration of CCl_4_ and LPS in C57BL/6 mice. **(B)** Flow cytometry gating strategy for hepatic and circulating cells **(C)** Quantification of M-MDSC or PMN-MDSC cells in liver and blood of CCl_4_ controls and CCl_4_+LPS groups. **(D)** Immunofluorescence micrographs of tissue sections from FFPE liver tissue. AF488=green depicts CD11b expression, AF647=red depicts Gr-1 expression, DAPI=blue depicts cell nuclei, scalebars=20µm. **(E)** Quantification per 10HPF magnification (400x) of CD11b+Gr-1+ cells **(F)** Representative images of PSR stains or H&E stains of liver sections, scalebar=100µm. **(G)** Ishak Stage and Ishak Grade scoring of stains and CPA analysis of PSR stains. Barplots displaying all datapoints with mean ± SEM, statistical tests by unpaired t-tests, *p < 0.05, **p < 0.01, ***p < 0.001.

We next characterised a model of CCl_4_-induced liver fibrosis with intravenous administration of a sublethal dose of *E. coli* 24 hours prior to sacrifice (fig. 5A). Given that the identification of MDSC subsets using conventional markers has proven to be challenging in previous studies[31], we further identified M-MDSC and PMN-MDSC as CD45^+^CD11b^+^Ly6C^hi^Ly6G^-^ CD84^+^ and CD45^+^CD11b^+^Ly6C^low^Ly6G^+^CD84^+^, respectively (fig. 5B and fig. S3C) [8, 29]. In the presence of infection, flow cytometry analysis of liver non-parenchymal cells and blood cells revealed an expansion of both M-MDSC and PMN-MDSC subsets in the liver and circulation of CCl_4_-treated mice compared to infection controls (liver: M-MDSC 5.23-fold change p=0.0004, PMN-MDSC 2.41-fold change p=0.0209; blood: M-MDSC 2.38-fold change p=0.0414, PMN-MDSC 8.18-fold change p<0.0001) (fig. 5C). Immunofluorescence on liver sections confirmed the expansion of CD11b^+^Gr-1^+^ MDSC subsets in CCl_4_-treated mice with infection compared to infection controls (2.86-fold change p=0.0012) (fig. 5D and 5E). Liver fibrosis was absent in infection controls (olive oil + *E. coli*), while CCl_4_-treated mice with infection displayed fibrosis as well as profound tissue necroinflammation, indicated by a high Ishak score (fig. 5F and 5G). Similar to CCl_4_-treated mice without infection, plasma markers of CCl_4_-treated mice with infection indicated liver damage but no impairment in liver synthetic function (fig. 5H).

**Figure 5.**
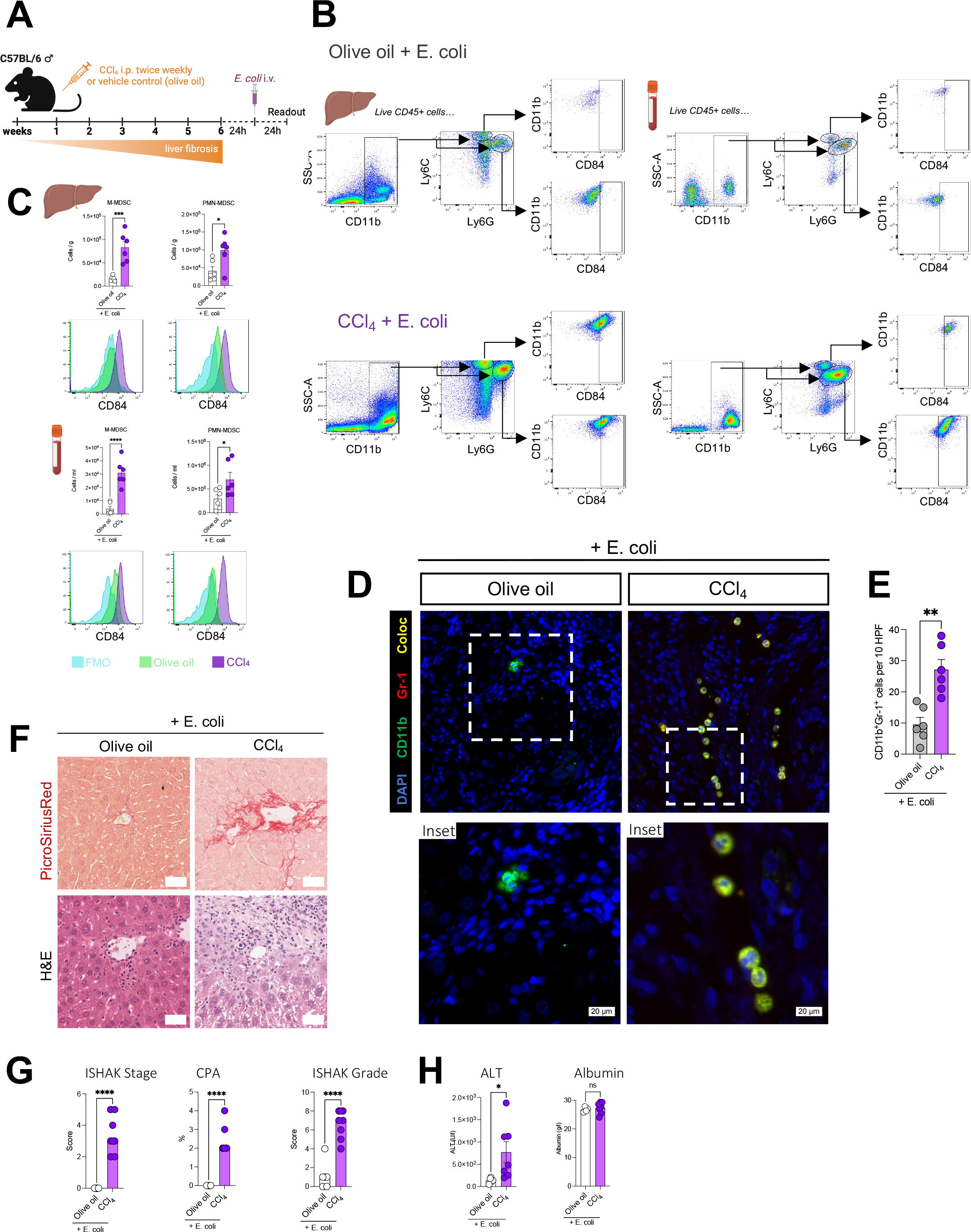
MDSC subsets expand in a model of CCl_4_-induced fibrosis with *E. coli* infection **(A)** Schedule of administration of CCl_4_ and i.v. *E.coli* administration in C56BL/6 mice. **(B)** Flow cytometry gating strategy for M-MDSC and PMN-MDSC in liver and blood in vehicle+E.coli controls and CCl_4_+E.coli groups **(C)** Quantification of M-MDSC or PMN-MDSC cells in liver and blood. CD84-BV786 expression in experimental groups and FMO. **(D)** Immunofluorescence micrographs of tissue sections from FFPE liver tissue from vehicle+E.coli and CCl_4_+E.coli groups. AF488=green depicts CD11b expression, AF647=red depicts Gr-1 expression, DAPI=blue depicts cell nuclei, scalebars=20µm. **(E)** Quantification per 10HPF magnification (400x) of CD11b+Gr-1+ cells **(F)** Representative images of PSR stains or H&E stains of liver sections, scalebar=100µm. **(G)** Ishak Stage and Ishak Grade scoring of stains and CPA analysis of PSR stains **(H)** Plasma levels of liver-related markers alanine aminotransferase and albumin. Barplots displaying all datapoints with mean ± SEM, statistical tests by unpaired t-tests (for n=2 groups) or one-way ordinary ANOVA with multiple comparisons (for n>2 groups), *p < 0.05, **p < 0.01, ***p < 0.001, ****p < 0.0001.

### TLR3 agonism limits expansion of hepatic MDSC and reduces fibrosis in a model of liver fibrosis with infection

Given that stimulation with poly(I:C) reduced the fraction of M-MDSC and improved innate immune function of these cells in patients with cirrhosis *ex vivo*, we explored the safety and potential benefit of poly(I:C) administration in a murine model of CCl_4_-induced liver fibrosis. We administered a low (1 mg/Kg) and a high dosage (4 mg/Kg) of poly(I:C) in CCl_4_-treated mice, 4 times over the course of the final week before sacrifice (fig. 6A).

**Figure 6.**
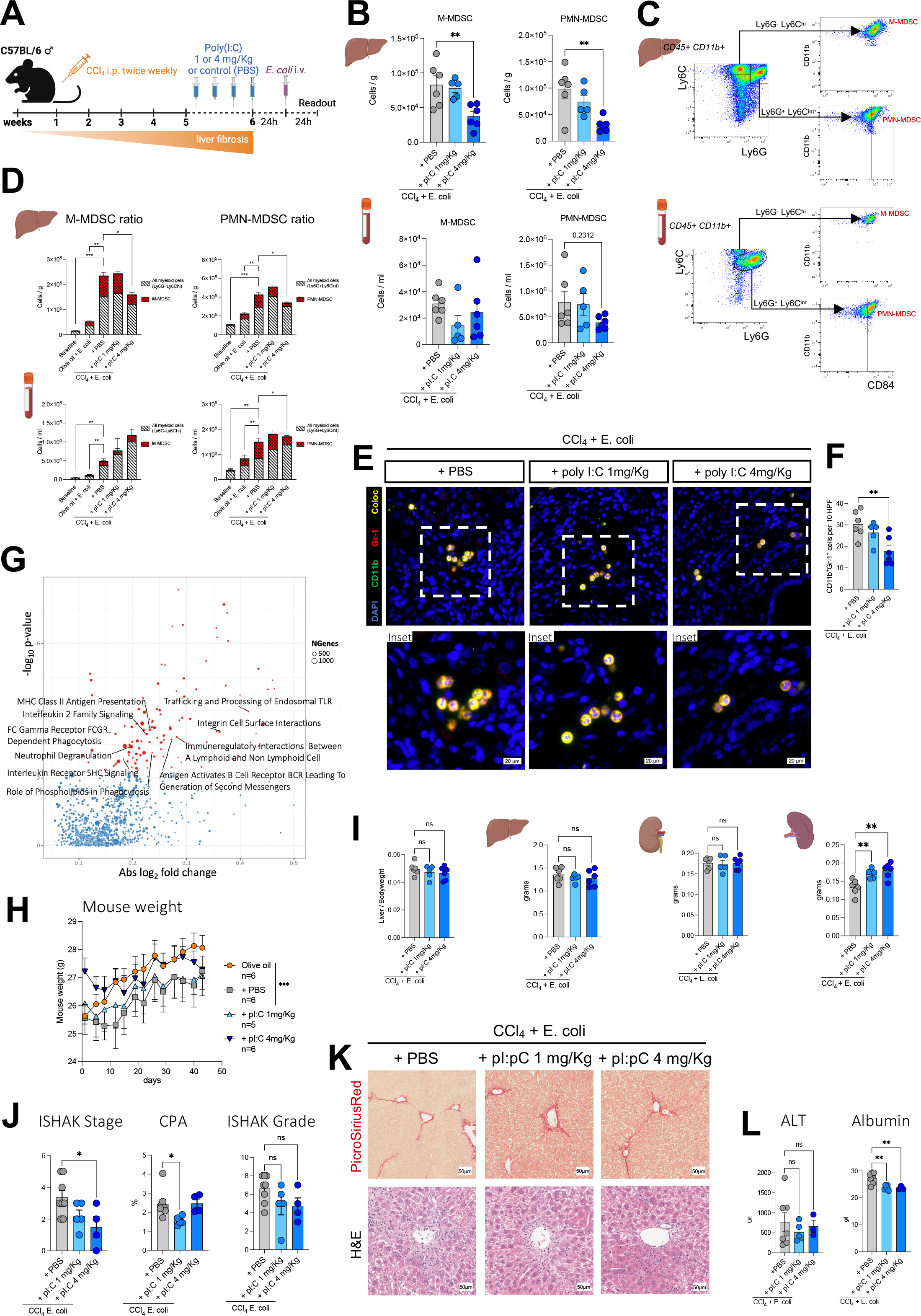
TLR3 agonism limits expansion of hepatic MDSC and reduces fibrosis in a model of liver fibrosis with infection. (A) Schedule of administration of CCl_4_ and i.v. E.coli, and poly(I:C) therapy (1 mg/Kg or 4 mg/Kg dosages) in C56BL/6 mice. **(B)** Quantification of M-MDSC or PMN-MDSC cells in liver and blood. **(C)** Flow cytometry gating strategy for M- MDSC and PMN-MDSC with parent myeloid populations Ly6G^-^Ly6C^hi^ and Ly6G^+^Ly6C^int^. **(D)** Stacked bar graphs representing the cell number ratios in the liver and blood of M-MDSC or PMN-MDSC to total myeloid cells across all experimental groups **(E)** Immunofluorescence micrographs of tissue sections from FFPE liver tissue. AF488=green depicts CD11b expression, AF647=red depicts Gr-1 expression, DAPI=blue depicts cell nuclei, scalebars=20µm. **(F)** Quantification per 10HPF magnification (400x) of CD11b+Gr-1+ cells. **(G)** Pathway enrichment analysis using the gene sets from the Reactome database (significant at an FDR of 5% in red) of BulkRNAseq samples from liver tissue of CCl_4_-treated mice with poly(I:C) stimulation vs CCl_4_ controls. **(H)** Bodyweight curves over 6 weeks, statistical comparison by two-Way ANOVA mixed model. **(I)** Liver-to-bodyweight ratio and organ weight of liver, kidney and spleen **(J)** Ishak Stage and Ishak Grade scoring of stains and CPA analysis of PSR stains. **(K)** Representative images of PSR stains or H&E stains of liver sections, scalebar=100µm. **(L)** Plasma levels of liver-related markers ALT and albumin. Barplots displaying all datapoints with mean ± SEM, statistical tests by one-way ordinary ANOVA with multiple comparisons, *p < 0.05, **p < 0.01, ***p < 0.001

Notably, a high dosage of poly(I:C) reduced the expansion of hepatic M-MDSC and PMN-MDSC, both in terms of absolute cell numbers as well as frequency relative to total CD45+ cells (M-MDSC –54.89% p=0.0074, PMN-MDSC –69.06% p=0.0036) (fig. 6B and S4A). In the circulation, MDSC subsets were numerically reduced following TLR3 agonism, although the difference was not statistically significant (fig. 6B and S6B). In addition, the ratio of M-MDSC to Ly6G^-^Ly6C^hi^ myeloid cells as well as the ratio of PMN-MDSC to Ly6G^+^Ly6C^int^ myeloid cells were reduced in the liver (fig. 6C and 6D), indicating that TLR3 agonism reduced the expansion of MDSC subsets in a specific manner. Overall, poly(I:C) therapy had no effect on frequency of CD45+ immune cells in both the liver and circulation, indicating no worsening of inflammation in these compartments compared to CCl4 with infection controls (fig. S4B). We confirmed our flow cytometry data from the liver by performing immunofluorescence for CD11b^+^Gr-1^+^ MDSC in liver sections, which indicated that a high dosage of poly(I:C) therapy reduced the expansion of MDSC compared to CCl_4_ with infection controls (–41.21% p=0.0068) (6E and 6F). In the circulation, poly(I:C) therapy reduced the ratio of PMN-MDSC to Ly6G^+^Ly6C^int^ myeloid cells, while the ratio of M-MDSC to Ly6G^-^Ly6C^hi^ myeloid cells remained unchanged (fig. 6C and 6D). Overall poly(I:C) stimulation had no effect on the numbers of hepatic lymphocyte subpopulations (fig. S4C). Bulk RNA seq of murine liver tissue followed by pathway enrichment analysis (Reactome database) revealed a variety of affected pathways in poly(I:C)-treated mice compared to CCl_4_ controls, including upregulation of pathways related to innate immunity (MHC II presentation, neutrophil degranulation, FCg receptor-dependent phagocytosis), interleukin-2 signaling and trafficking/processing of endosomal TLR (fig. 6G).

Bodyweight of poly(I:C)-treated mice was not affected in the final week of therapy compared to CCl_4_ controls (fig. 6H). The liver to bodyweight ratio as well as the liver and kidney weights of poly(I:C)-treated animals were comparable to CCl_4_ controls. However, poly(I:C) therapy increased spleen weight compared to fibrotic controls (6I). Remarkably, Ishak Stage scoring and CPA analysis indicated that poly(I:C) therapy reduced liver fibrosis. In addition, Ishak grade scores of hepatic necroinflammation were numerically reduced for both dosages of poly(I:C), although these changes did not reach statistical significance (fig. 6J and 6K). Plasma levels of ALT were comparable indicating no effect on liver damage following poly(I:C) therapy, while albumin levels were reduced following both dosages (fig. 6L).

Taken together, these data indicated that a high dose of poly(I:C) selectively reduced the expansion of M-MDSC and PMN-MDSC in the liver tissue of CCl_4_-treated mice with infection, without exacerbating inflammation in the liver and circulation. In addition, a high dosage of poly(I:C) selectively reduced circulating PMN-MDSC. Notably, poly(I:C) therapy reduced hepatic fibrosis compared to fibrotic controls. Overall, these data underline the potential beneficial effect and liver-related safety of poly(I:C) therapy *in vivo* in a model of CCl_4_-induced fibrosis.

## Discussion

In this study we detailed the expansion and clinical significance of MDSC subsets in both the circulation and liver of patients with cirrhosis. While in a previous study we described an accumulation of circulating monocytic MDSC in advanced stages of cirrhosis and ACLF[16], here we revealed their accumulation also in the liver. Moreover, we also observed the accumulation of polymorphonuclear MDSC in the liver, which may contribute to further dampen immune responses. As in the circulation, the expansion of MDSC subsets in the liver was strongly correlated with indices of disease severity, clinical variables, infectious complications and reduced transplant-free survival, thereby highlighting their biological relevance. In addition, we detailed the expansion of immunosuppressive MDSC subsets in both the circulation and liver in murine models of chronic liver injury. Subsequently, we used a model of CCl_4_-induced liver fibrosis with bacterial infection to investigate the safety and therapeutic efficacy of poly(I:C) therapy targeting the expansion of MDSC *in vivo*.

A large number of clinical studies in the field of sepsis indicated that M-MDSC may represent not only attractive biomarkers but also potential therapeutic targets to improve patient outcome[15]. Previously, we had reported the predominance of circulating M-MDSC in patients with AD/ACLF, but also to a lower extent in patients with compensated cirrhosis[16]. These cells were characterised by attenuated cytokine responses to TLR stimulation and impaired phagocytosis of pathogens, thereby representing a cellular population that contributes to immuneparesis and infection susceptibility in cirrhosis. Moreover, M-MDSC in the blood strongly indicated reduced survival[16, 28]. Here, the observation that M-MDSC expanded not only in the circulation, but also in the liver and the peritoneal cavity along with cirrhosis progression, is intriguing. Recently, in addition to other malignancies, M-MDSC were identified in the tumour microenvironment of HCC lesions[32–34]. While it is assumed that their biological function may be to control chronic tissue inflammation[35], their origin in the context of liver cirrhosis remains unknown. Exposure to the circulating blood milieu of patients with cirrhosis, specifically PAMPs such as TLR ligands and cytokines, led to the differentiation of M-MDSC *in vitro*[16]. In fact, in a murine model, we noted a further expansion of M-MDSC in the liver after acute stimulation with LPS compared to fibrosis controls. However, even in the absence of PAMP stimulation, M-MDSC were observed in the liver and the blood in a murine model of CCl_4_-induced liver fibrosis after 6 weeks. These findings may suggest that M-MDSC differentiate in the circulatory milieu, as previously shown, and subsequently infiltrate tissues such as the liver[36]. In parallel, we noted a reduction of distinct liver-resident Kupffer cells in our murine models *in vivo*, in accordance with previous observations in a CCl_4_-induced liver fibrosis model[37]. While a study on patients with chronic hepatitis B revealed that HBe-Ag led to the expansion of hepatic M-MDSC and to a dampening of CD8+ T cell function[38], these findings do not explain M-MDSC emergence in patients with cirrhosis of non-viral aetiology. The expansion of M-MDSC may be due to changes within the liver and peritoneal milieu themselves, such as cirrhosis-associated bacterial translocation[39] which was previously postulated for their emergence in HCC[32]. In addition, other tissue-specific processes may account for M-MDSC expansion, including hepatic stellate cells that lead to an accumulation of M-MDSC in the liver both through recruitment (via COX2-PGE2-EP4 or p38 MAPK signaling) and through a direct-contact mechanism with M-MDSC[40]. However, in our study, bone marrow tissue obtained from a cirrhotic patient prior to bone marrow transplantation displayed an expansion of M-MDSC compared to a bone marrow control, which is in accordance with a previous study in ACLF patients with underlying HBV infection[41].

PMN-MDSC were also identified throughout numerous clinical studies in patients with sepsis. These cells were heavily linked to poor outcomes and represent a promising therapeutic target[15]. Interestingly, in the present study we noted an expansion of PMN-MDSC both in the liver of patients with cirrhosis and in our mouse models of chronic liver injury, which may further contribute to the state of immuneparesis in cirrhosis. PMN-MDSC in the tumour microenvironment of HCC have been previously reported to facilitate HCC development, and therapies targeting PMN-MDSC are being investigated[33, 34, 42–45]. However, to our knowledge, our study is the first to show an expansion of PMN-MDSC in tumour-free cirrhotic liver tissue, which served as an indicator of adverse prognosis. Albeit, a previous study had highlighted an accumulation of granulocytic MDSC in the blood and liver of patients with ACLF with underlying HBV infection, which were associated with impaired immune responses (including suppression of T cell proliferation, cytokine secretion, phagocytosis) and indicated infection and death[41]. In addition, both *ex vivo* and *in vivo* data from a mouse model of Concanavalin-induced ACLF indicated that the generation of PMN-MDSC was dependent on plasma milieu component TNF-α, although PMN-MDSC also expanded in the bone marrow[41]. In another study, mouse models for primary sclerosing cholangitis and colitis displayed gut barrier dysfunction and bacterial translocation, which led to an accumulation of CXCR2+ PMN-MDSC in the liver through a TLR4-dependent mechanism[46]. Accordingly, in our CCl_4_-induced liver fibrosis model we noted further expansion of PMN-MDSC both in the blood and liver following an acute challenge with LPS, suggesting a role for bacterial exposure in the process of PMN-MDSC accumulation. However, we also observed PMN-MDSC expansion in the bone marrow of a patient with decompensated cirrhosis. Thus, further studies on the ontogeny and migration patterns of both M-MDSC and PMN-MDSC in cirrhosis are needed.

Within the field of MDSC biology, a significant challenge stems from the broad categorisation of MDSCs, which encompasses a diverse array of cells with immunosuppressive capabilities, notably affecting both adaptive and, as recent studies including this one have shown, innate immunity[16]. In fact, MDSC can originate from different developmental stages, appear across various disease contexts, employ distinct immunosuppressive tactics and showcase unique phenotypes[8, 36]. Thus, caution is needed when comparing MDSCs across different studies, as they may likely represent disparate cell populations. While in this study we successfully identified MDSC subsets both in patients with cirrhosis and in murine models of chronic liver injury, it remains challenging to assert that these cells are indeed analogous, given the broad differences in species-specific phenotypical markers, disease states and experimental protocols. Furthermore, even though further studies into the ontogeny of MDSCs in murine models (through for example fate mapping experiments) may provide exhaustive answers, extending such detailed investigations to human subjects with cirrhosis presents substantial technical hurdles.

Given that the accumulation of MDSC subsets in cirrhosis is linked to immuneparesis and increased infection susceptibility, we pondered whether the generation and expansion of MDSC may be prevented with immunotherapy. In the context of other pathologies, previous *in vivo* studies in mice demonstrated a variety of effects of poly(I:C) therapy not only on innate immunity but also on MDSC. Briefly, poly(I:C) promoted phagocytic activity of bacteria by peritoneal macrophages[17] and enhanced infection clearance and survival in a secondary pneumonia model[18]. In addition, poly(I:C) therapy reduced MDSC frequency in the tumour microenvironment (TME) of liver metastases[19–21]. Poly(I:C) therapy reduced the suppressive capacity of both M-MDSC and PMN-MDSC on T cells in the TME of a murine orthotopic pancreatic cancer model and this immunosuppression was dependent on IFNAR1 signaling[47]. Moreover, poly(I:C) therapy decreased MDSC numbers in the peripheral blood and spleen of mice bearing subcutaneous colon or orthotopic breast cancer xenografts[48]. Importantly, we previously demonstrated that stimulation with poly(I:C) *ex vivo* in patients with cirrhosis reduced M-MDSC frequency and reversed the impaired innate immunity of these cells by restoring the phagocytosis capacity of *E. coli*[16]. In the present study, poly(I:C) stimulation *ex vivo* of M-MDSC from patients with cirrhosis increased HLA-DR expression on these cells, reduced the frequency M-MDSC, reduced the expression of CD84 – a novel MDSC-specific marker[27, 29] – and enhanced the phagocytic capacity of both *E. coli* and *S. aureus* by these cells. In addition, *in vivo* stimulation with poly(I:C) in our murine model of CCl_4_-induced liver fibrosis with infection reduced the accumulation of both M-MDSC and PMN-MDSC subsets in the liver in a dose-dependent manner, while MDSC subsets were also numerically reduced in the blood. These findings underline the efficacy of poly(I:C) therapy at reducing the accumulation of pathological hepatic MDSC subsets in the context of chronic liver injury *in vivo*.

Besides the immunomodulatory effects of poly(I:C), multiple studies indicated that poly(I:C) administration attenuates hepatic fibrosis in murine models, including *Clonorchis sinensis* infection and *Schistosoma japonicum* egg-induced liver fibrosis models[24, 26]. In addition, another study showed that either poly(I:C) or IFN-γ inhibited CCl_4_-induced liver fibrosis in mice, and that fibrosis reduction was dependent on STAT1 signaling[25]. In our murine model, poly(I:C) therapy also reduced CCl_4_-induced hepatic fibrosis. Importantly, we did not observe significant off-target side effects due to poly(I:C) stimulation, with the exception of marginally reduced serum albumin levels and increased spleen weight compared to controls. The lower serum albumin levels might not be necessarily due to reduced liver synthetic function, but could be the result of processes associated with poly(I:C)-induced gut inflammation such as protein-losing enteropathy and malabsorption[49]. In support of its safety profile, poly(I:C) as a vaccine adjuvant was generally well tolerated in a phase II clinical trial[50]. Apart from this clinical trial, currently the use of poly-ICLC (i.e. an immunostimulant composed of poly(I:C) mixed with stabilizers carboxymethylcellulose and polylysine[51]) is currently preferred over the use of poly(I:C) in phase II clinical trials in cancer, due to its superior capacity to bolster both innate and adaptive immunity against cancer cells[52].

Given that poly(I:C) and TLR3 activation are known to play pivotal roles in a variety of physiological and pathological contexts (most notably antiviral responses but also inflammatory responses, anticancer immunity and autoimmunity) through the production of Type I IFNs (IFN-α and IFN-β)[53], it would be interesting to investigate whether the poly(I:C)- induced effects that we observed in the present study could be replicated by stimulation with IFN-α and/or IFN-β directly.

In conclusion, our study highlights the biological significance of MDSC expansion during progression of cirrhosis, which was independent of underlying aetiology, was associated with dampened innate immune responses, susceptibility to infections and reduced transplant-free survival. Poly(I:C) therapy was evaluated both *ex vivo* in patients with cirrhosis and *in vivo* in our murine model and represents a promising immunomodulatory agent to prevent MDSC accumulation and reverse impaired innate immune responses in cirrhosis. Thus, we propose its further evaluation in human studies *in vivo* to assess its preventive character on infectious complications in cirrhosis.

## Abbreviations

MDSC: myeloid-derived suppressor cells, M-MDSC: monoytic myeloid-derived suppressor cells, PMN-MDSC: polymorphonuclear myeloid-derived suppressor cells, poly(I:C): polyinosinic:polycytidylic acid, CCl_4_: carbon tetrachloride LPS: lipopolysaccharide, AD: acute decompensation, ACLF: acute-on-chronic liver failure, MELD: model for end-stage liver disease, INR: international normalised ratio, ALT: alanine transaminase, PSR: picrosirius red, KC: Kupffer cells, TLR: toll-like receptor, PAMP: pathogen-associated molecular pattern, HCC: hepatocellular carcinoma, HBV: hepatitis B virus, TNF-α: tumour necrosis factor alpha, TME: tumour microenvironment, PBMC: peripheral blood mononuclear cells, MFI: mean fluorescence intensity, HPF: high-power field, FMO: fluorescence minus one.

## Material and Methods

### Patients and sampling

We investigated 22 patients with cirrhosis that were recruited at the University Hospital Basel and the Cantonal Hospital St. Gallen, Switzerland, between January 2016 and December 2017. All patients provided written informed consent and were categorised based on Child-Pugh scores (Child-Pugh A n = 8, Child-Pugh B n=7, Child-Pugh n=7). Additionally, we included 4 control subjects without liver disease. All liver sections were obtained from liver biopsies. Exclusion criteria included age under 18, infection with human immunodeficiency virus, immunosuppressive therapy and malignancy.

### Clinical, haematological and biochemical data

Data related to clinical variables, full blood counts as well as liver and renal function tests were prospectively entered into a database. Child-Pugh and MELD disease severity scores were calculated, survival, transplantation status, infections and development of HCC were recorded.

### Flow cytometry-based phenotyping of monocytes

Phenotyping of circulating monocytes was performed with flow cytometry as previously described[16]. Flow cytometric data was acquired on a BD LSRFortessa Cell Analyzer (BD Biosciences) and data was analysed using FlowJo version 10.9. Results are depicted either as percentage of positive cells or by mean fluorescence intensity. For further details on flow cytometry antibodies see supplementary information.

### *Ex vivo* phagocytosis assay in PBMC

Phagocytosis capacity was assessed for 500’000 PBMCs per well in a 24-well plate in 500µl medium (RPMI with 10% FBS, 1% penicillin-streptomycin) supplemented with 20% of human AB serum (Sigma-Aldrich Cat.# H3667). PHrodo E. coli BioParticles and pHrodo S.aureus BioParticles (ThermoFisher Cat.# P35361, A10010) were incubated for 60min at 37°C, 5% CO_2_. Cells were analysed with flow cytometry as detailed above.

### Carbon tetrachloride-induced liver fibrosis model

Wild-type C57BL/6 mice aged between 8 to 10 weeks were purchased from Charles River Laboratories and were administered CCl_4_ 0.4 mL/Kg (Sigma-Aldrich), which was diluted in olive oil (Sigma-Aldrich) at a ratio of 1:3. Intraperitoneal (i.p.) injections were performed bi- weekly over a total of 6 weeks. The CCl_4_-only model and the CCl_4_ with bacterial challenge model included 13 CCl_4_ injections (fig. 3, 5, 6), the last injection was 24h prior to sacrifice. The LPS challenge model (fig. 4) included 12 CCl_4_ injections, the last CCl_4_ injection was 96h prior to sacrifice. Following sacrifice, livers were perfused with PBS and liver tissue was stored for flow cytometry analysis, RNA isolation, as well as formalin fixation and paraffin embedding for further analyses.

### *In vivo* treatments

A group of CCl4-treated mice was intraperitoneally administered 100 μg of LPS per mouse, 72h after the final CCl_4_ injection and 24h prior to sacrifice. In the presence of bacterial challenge with intravenous administration of *E.coli*, two groups of CCl_4_-treated mice were intraperitoneally administered poly(I:C) at dosages of either 1 mg/Kg or 4 mg/Kg as well as a control group (PBS i.p.).

### Bacterial challenge

*E. coli* (25922GFP, American Type Culture Collection [ATCC]) were cultured in Tryptic Soy Agar/Broth (MilliporeSigma). Bacteria was administered intravenously at a nonlethal dose of 5 x 10^7^ *E. coli* per 20 grams of body weight. Disease severity was assessed using a murine sepsis scoring system. Monitoring of animal health was performed by 2 investigators independently every 2 hours for the first 8 hours, which included evaluations of appearance, level of consciousness, activity, response to stimuli, eye health as well as respiration rate and quality[54].

### Flow cytometry-based phenotyping of murine hepatic and blood cells

Murine hepatic non-parenchymal cells were isolated as previously described[55]. Myeloid and lymphoid cells from the liver as well as blood myeloid cells were phenotypically characterised by flow cytometry. Isolated cells were treated for 5 min with TruStain FcX^TM^ (anti-mouse CD16/CD32) (cat. #101320, BioLegend) prior to staining, then cells were stained with monoclonal antibodies for 30 min at 4°C in Brilliant Stain Buffer (BioLegend, cat. #563794). See supplementary information for details on flow cytometry antibodies. Cells were analysed using flow cytometry analysis as detailed above. Frequency of cell subsets was normalized either to gram of liver tissue or to millilitre of blood using 123count^TM^ eBeads Counting Beads (ThermoFisher, cat. #01-1234-42).

### T cell proliferation assay

Liver non-parenchymal cells were isolated from healthy controls and CCl_4_-treated mice, either M-MDSC or PMN-MDSC were FACS sorted (BD FACS Melody sorter) and co-cultured in a 1:1 ratio with CD4+ T cells isolated from the livers of healthy control mice, using the MojoSort magnet with MojoSort Mouse CD4 Naïve T Cell Isolation Kit (Cat.#480040 BioLegend). T cell stimulation was induced with anti-CD3ε and anti-CD28 beads (T Cell Activation/Expansion Kit mouse, Cat.#130-093-627, Miltenyi Biotec) according to manufacturer’s instructions. Cell doublings were tracked with a CellTrace CFSE Cell Proliferation Kit (Cat.#C34554 Invitrogen). Cells were co-cultured in RPMI 1640 medium (10% fetal bovine serum, 1% Pen/Strep), cell proliferation was assessed after 4 days. Cells were analysed with flow cytometry as detailed above. See supplementary information for details on antibodies.

### Flow cytometry-based phenotyping of peritoneal macrophages

Peritoneal macrophages (pMacs) were isolated from ascites by centrifugation following paracentesis in patients with cirrhosis presenting ascites. The pMacs were stained and analysed by flow cytometry as previously described[56].

### Fluorescent immunohistochemistry and microscopy

All sections from FFPE tissues were 4 μm-thick. Prior to staining, tissue sections were processed as previously described[56] (see supplementary material and methods for further details). Multiplexed immunofluorescence on FFPE tissue was performed to assess CD11b and Ly-6G/Ly-6C (Gr-1) expression in murine liver sections, as well as CD14, CD15 and CD84 in sections from human liver biopsies. Slides were mounted using Fluoromount-G^TM^ containing DAPI (ThermoFisher, cat. #00-4959-52).

### Image acquisition and analysis

Micrographs of fluorescent stains were acquired using a Nikon TI2 widefield microscope equipped with a Photometrics Prime 95B camera (Teledyne Photometrics). Quantitative morphometry was performed on 10 HPF (400x magnification) per slide assessing the following cell subsets: CD11b^+^Gr-1^+^ (mouse), CD14^+^CD84^+^ and CD15^+^CD84^+^ (human).

### Histopathalogical analysis of PSR and H&E stains

Histological assessment was performed by a trained histopathologist (C.E.) on H&E and PSR stains of FFPE liver tissue sections from mice across all murine experimental conditions. Samples were evaluated according to Ishak grading and staging systems[57]. Histological assessment was performed according to Ishak Staging and Ishak Grading, including confluent necrosis, focal lytic necrosis, portal and periportal inflammation, steatosis grade (macrovesicular), hepatocyte ballooning, hypertrophy, glycogenated nuclei and apoptotic bodies. CPA analysis was performed as PSR-positive area compared to total surface area on PSR-stains of FFPE liver sections using Fiji (v. 2.9.0).

### Quantification of liver-related plasma biomarkers

Blood was collected and plasma was isolated from all murine experimental groups. Plasma levels of albumin were assessed with the bromocresol green method, plasma alanine aminotransferase (ALT) levels were quantitatively determined using the modified method without pyridoxal phosphate, on a AU680 analyser (clinical chemistry tests, Mary Lyon Centre at MRC Harwell).

### Bulk RNA sequencing

Liver tissue was harvested from healthy controls, CCl_4_-treated controls and CCl_4_-treated mice with poly(I:C) therapy (1 mg/Kg) and stored in RNAlater stabilization solution (ThermoFisher, cat.# AM7020). Tissue homogenisation was performed using a Fast-Prep 24-5G instrument (MP Biomedicals), total RNA was extracted using the RNeasy Plus Mini Kit (Qiagen, cat.#74134).

Library preparation was performed starting from 200ng total RNA of each sample using the TruSeq Stranded mRNA Library Kit (Illumina, cat. # 20020595) and the TruSeq RNA UD Indexes (Illumina, cat. # 20022371). Gene set enrichment analysis was performed with the function *camera* from the edgeR package[58] (using the default parameter value of 0.01 for the correlations of genes within gene sets) using gene sets from the mouse collections of the MSigDB Molecular Signatures Database[59] (version 2023.2). We notably focused on the curated gene sets (M2 collection) and its Reactome subset of canonical pathways. We filtered out sets containing less than 10 genes, and gene sets with an FDR lower than 5% were considered significant. For further details, including bioinformatic data analysis, see supplementary information.

### Statistical analysis

Data were presented and analysed using GraphPad Prism 10 (GraphPad Software). Data are presented as mean ± SEM. For data that followed a normal distribution, unpaired t tests and One-Way ANOVA with Holm-Šidák multiple comparisons were used for statistical analysis of differences between 2 groups and among n > 2 groups, respectively. For non-normally distributed data, Mann-Whitney *U* nonparametric and Kruskal-Wallis tests were used for statistical analysis of differences between 2 groups and among n > 2 groups, respectively. For the statistical analysis of differences among n > 2 groups with 2 independent variables a 2- way ANOVA with Tukey’s multiple comparisons test was used. To assess correlations with clinical parameters, a Spearman’s nonparametric correlation test was used. P values below 0.05 were considered statistically significant.

### Study approval

All subjects provided written informed consent. The human study was approved by local ethics committees (EKSG 15/074/EKNZ 2015-308, UK:12/LO/0167). Animal experimental protocols were in accordance with UK Home Office regulations and were approved by Imperial College London (PPL number P8999BD42).

### Data availability statement

Data are available in a public, open access repository. GEO Accession GSE260935, https://www.ncbi.nlm.nih.gov/geo/query/acc.cgi?acc=GSE260935.

## Acknowledgments

The authors thank all patients who consented to participate in this study and all staff at the University Hospital Basel and the Cantonal Hospital St. Gallen involved in these patients’ care. The authors also thank the Department of Biomedicine of the University of Basel for infrastructural support, in particular the core facilities including the Bioinformatics Core, Flow Cytometry Core, Histology Core and Microscopy Core facilities. Also, the authors are grateful to the Gastroenterology Research Group, led by Prof. J. Niess, Department of Biomedicine, for methodological and conceptual advice. Calculations were performed at sciCORE (http://scicore.unibas.ch/) scientific computing center at University of Basel. Figures were created with BioRender.com.

