## Supplemental Information for "Targeting the expansion of myeloid-derived suppressor cells in liver cirrhosis"

### Supplementary information

#### Materials and methods

##### Flow cytometry antibodies

*Human:* The following flow cytometry antibodies were used to phenotypically characterise circulating monocytes isolated from patients with cirrhosis: BioLegend: CD14 PE-Cy7 (#301814), CD15 APC (#301908), HLA-DR PerCP Cy5.5 (#307630), CD3 APC/Fire750 (#300470), CD19 APC/Fire750 (#302258), CD56 APC/Fire750 (#362554). TLR3 PE (eBioscience, #12-9039-82), CD84 BV421 (BD Biosciences). Cell viability was assessed with a fixable viability dye eFluor 455UV (eBioscience #65-0868-18).

*Mouse:* The following flow cytometry antibodies were used to phenotypically characterise liver non-parenchymal cells: BioLegend: Zombie Live Dead (#423101), F4/80 BV421 (#123137), CD45-BV650 (#103151), CD11b BV711 (#101242), CD64 PerCP-Cy5.5 (#139308), Ly6c PE-Cy7 (#128018); Ly6G BV605 (#127639), CD8 BV711 (#100759), CD3 BV785 (#100232), CD19 PerCP (#115532), NK1.1 PE-Cy7 (#108714), CD4 AF647 (#100424), TCR $\beta$ APC/Cy7 (#109220); eBioscience: Tim4 AF488 (#53-5866-82), Mertk AF700 (#56-5751-82), MHCII APC-eFluor780 (#47-5321-82); BD Biosciences: CD84 BV785 (#749562); R&D: Tim-4 AF700 (#FAB2826N). The following antibodies were used for the mixed lymphocyte reaction: CD4 BV786 (BD Biosciences #563331), CD3 PE-Dazzle 594 (BioLegend, #100246).

##### Fluorescent immunohistochemistry
Murine liver and extrahepatic organs were fixed in 10% paraformaldehyde and were embedded in paraffin in the Histology Service of the Inflammation Repair and Development section of Imperial College. FFPE tissue sections were dewaxed in xylene, rehydrated in decreasing concentrations of ethanol. This was followed by heat-induced epitope retrieval at 95°C for 20 min in citrate buffer (pH 6) and a 1-hour incubation in 5% skim milk in PBS at room temperature (RT) for blocking of non-specific binding. Sections were incubated with primary antibodies overnight at 4°C, while the incubation with the secondary antibodies was for 1 hour at RT.
The following primary antibodies were used: CD11b (rabbit anti-mouse, Abcam, cat. #ab184308), Ly-6G/Ly-6C (Gr-1) (rat anti-mouse, R&D, cat. #MAB1037-SP), CD14 (mouse anti-human, Abcam, cat. #ab181470), CD15 (mouse anti-human, BioLegend, cat. #301902) and CD84 (rabbit anti-human, Abcam, cat. #131256). The following secondary antibodies were used: goat anti-rat AF488 (ThermoFisher, cat. #A-11006), donkey anti-rabbit AF647 (Abcam, cat. #ab150075), and goat anti-mouse AF488 (Abcam, cat. #ab150113).

##### Bulk RNA sequencing
Following total RNA extraction from murine liver tissue, the purity of each RNA was assessed using a NanoDrop One spectrophotometer (ThermoFisher Scientific). Samples were quantified by Fluorometry on a Qubit Flex Fluorometer (ThermoFisher Scientific) using the Qubit RNA High Sensitivity (ThermoFisher Scientific, cat. # Q32855). Their integrity was assessed on a TapeStation instrument (Agilent Technologies) using the High Sensitivity RNA ScreenTape (Agilent, cat. # 5067-5579). The average RINe of 8.7±0.5 showed high and consistant RNA integrity for all samples.
Libraries were quality-checked on the Fragment Analyzer instrument (Agilent Technologies) using the Standard Sensitivity NGS Fragment Analysis Kit (Agilent Technologies, cat. # DNF-473-0500). They were pooled to equal molarity. The pool was quantified by fluorometry and sequenced Paired-End 38 bases on NextSeq 500 (Illumina) using the NextSeq 500 High Output Kit 75-cycles (Illumina, cat. # FC-404-1005). The Flow cell was loaded at 2.0pM. 1% PhiX was included in the pool. On average per sample: 35±3 millions pass-filter

reads were collected. Data analysis was performed by the Bioinformatics Core Facility, Department of Biomedicine, University of Basel. Read quality was assessed with the FastQC tool (version 0.11.5). Reads were mapped to the mouse genome mm10 with STAR[1] (version 2.7.10a) with default parameters, except filtering out multimapping reads with more than 10 alignment locations (outFilterMultimapNmax=10) and filtering reads without evidence in the spliced junction table (outFilterType="BySJout"). The *featureCounts* function[2] from the Rsubread package (version 2.0.6) was used to count the number of reads (5' ends) overlapping with the exons of each gene (Ensembl release 102 genes) assuming an exon union model. All subsequent analyses were performed using the R software (version 4.3.1) and Bioconductor 3.18 packages[3]. The Bioconductor package edgeR (version 4.0.1) was used for differential gene expression analysis. Between samples normalization was performed using the TMM method. Genes with CPM values above 1 in at least 3 samples were retained for the differential expression analysis. The quasi-likelihood testing framework (edgeR function *glmQLFit* with optioin legacy=TRUE and function *glmQLFTest*) was used to perform pairwise gene expression comparisons between treatments. P-values were adjusted by controlling the false discovery rate (FDR; Benjamini-Hochberg method) and genes with an FDR lower than 5% were considered significant.

**Table S1 (excel)** Full results from the differential expression analysis performed on BulkRNAseq data from mouse liver tissue, including comparisons of healthy controls vs $CCl_4$-treated mice and $CCl_4$ controls vs $CCl_4$ with poly(I:C) therapy.

**Table S2 (excel)** Full gene set enrichment analysis of reactome pathways (using Reactome Database) performed on BulkRNAseq data from mouse liver tissue, including comparisons of healthy controls vs $CCl_4$-treated mice and $CCl_4$ controls vs $CCl_4$ with poly(I:C) therapy.

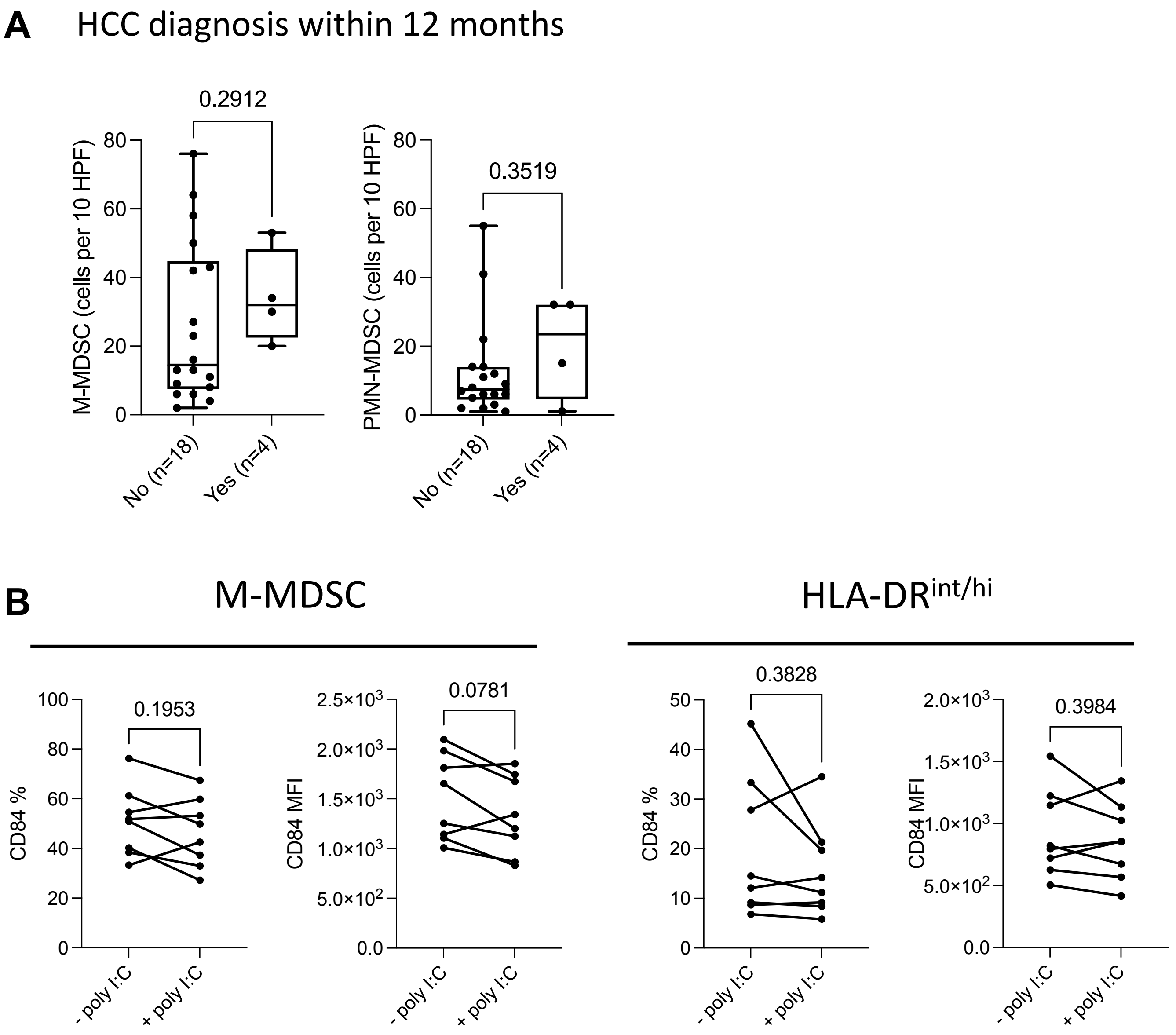
A
HCC diagnosis within 12 months
0.2912
M-MDSC (cells per 10 HPF)
0.3519
PMN-MDSC (cells per 10 HPF)
No (n=18)
Yes (n=4)
B
M-MDSC
HLA-DRint/hi
0.1953
CD84 %
0.0781
CD84 MFI
0.3828
0.3984
- poly I:C
+ poly I:C

**Figure S1. (A)** M-MDSC and PMN-MDSC numbers assessed by immunofluorescence of liver biopsies in relation to HCC diagnosis within a one-year timeframe **(B)** Percentage and MFI values of CD84 expression on cell subsets identified in fig. 2A with flow cytometry analysis. Statistical comparisons by Wilcoxon Tests (paired data).

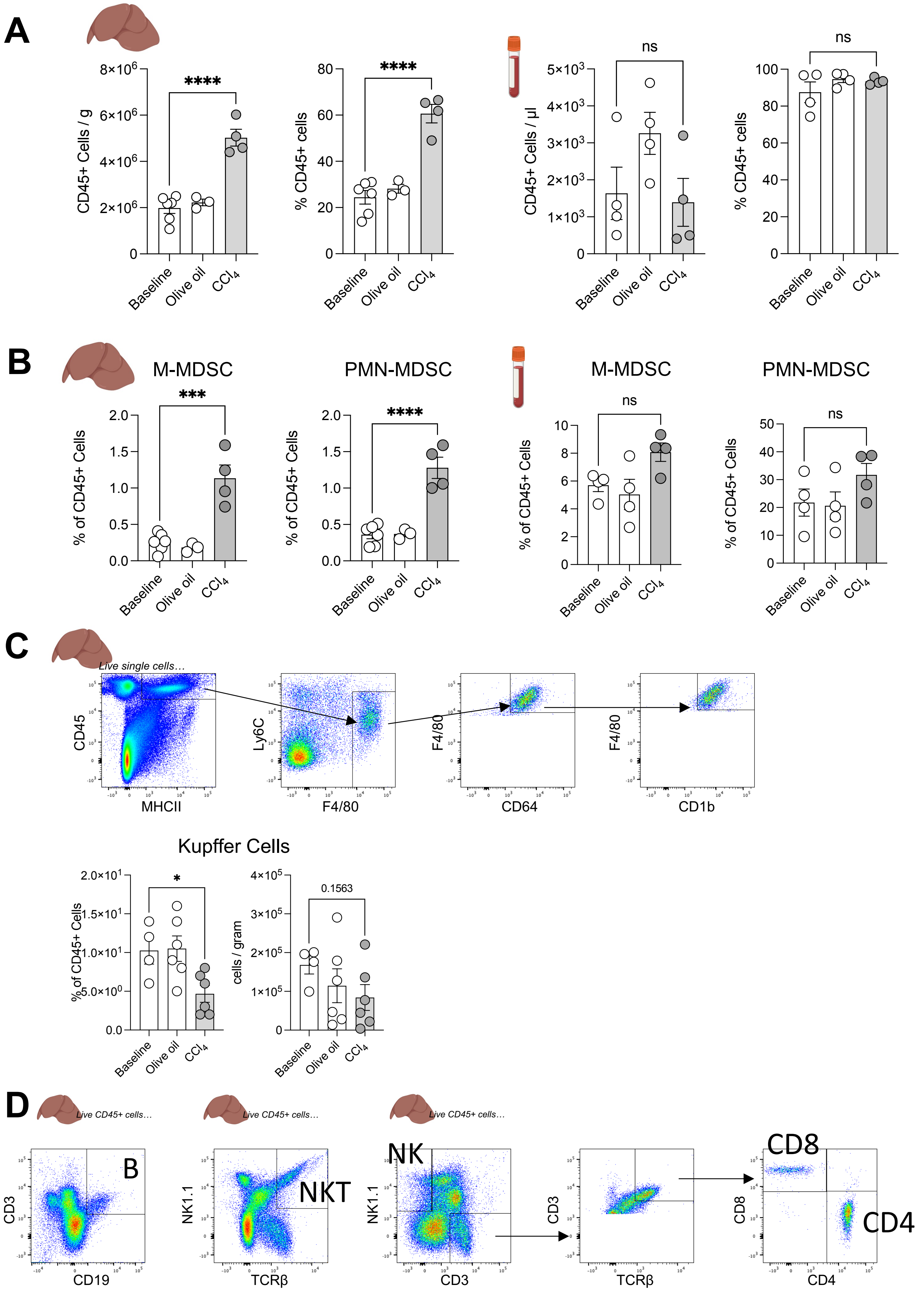

A
CD45+ Cells / g
% CD45+ cells
CD45+ Cells / µl
% CD45+ cells
****
****
ns
ns
Baseline
Olive oil
CCl4
B
M-MDSC
PMN-MDSC
M-MDSC
PMN-MDSC
% of CD45+ Cells
***
****
ns
ns
C
Live single cells…
CD45
MHCII
Ly6C
F4/80
F4/80
CD64
F4/80
CD1b
Kupffer Cells
% of CD45+ Cells
cells / gram
*
0.1563
D
Live CD45+ cells…
B
NKT
NK
CD8
CD4
CD3
CD19
NK1.1
TCRβ
NK1.1
CD3
CD3
TCRβ
CD8
CD4

**Figure S2. (A)** Cell numbers and percentages of CD45+ cells (of total live cells) in both liver (left) and blood (right) compartments identified with flow cytometry analysis, cell numbers were normalised either to gram of liver tissue or to µl of blood using flow cytometry counting beads. Groups include healthy controls, vehicle controls (olive oil) and $CCl_4$-treated mice. **(B)** Percentages of either M-MDSC or PMN-MDSC compared to total CD45+ cells in both liver (left) and blood (right) compartments identified with flow cytometry analysis. **(C)** Gating strategy for liver-resident Kupffer Cells (left), percentage of KC of total CD45+ immune cells and KC numbers per gram of liver tissue. **(D)** Flow cytometry gating strategy for hepatic lymphocyte subpopulations starting from live CD45+ cells isolated from murine livers Barplots displaying all datapoints with mean ± SEM, statistical tests by one-way ordinary ANOVA with multiple comparisons, ***$p < 0.001$, ****$p < 0.0001$.

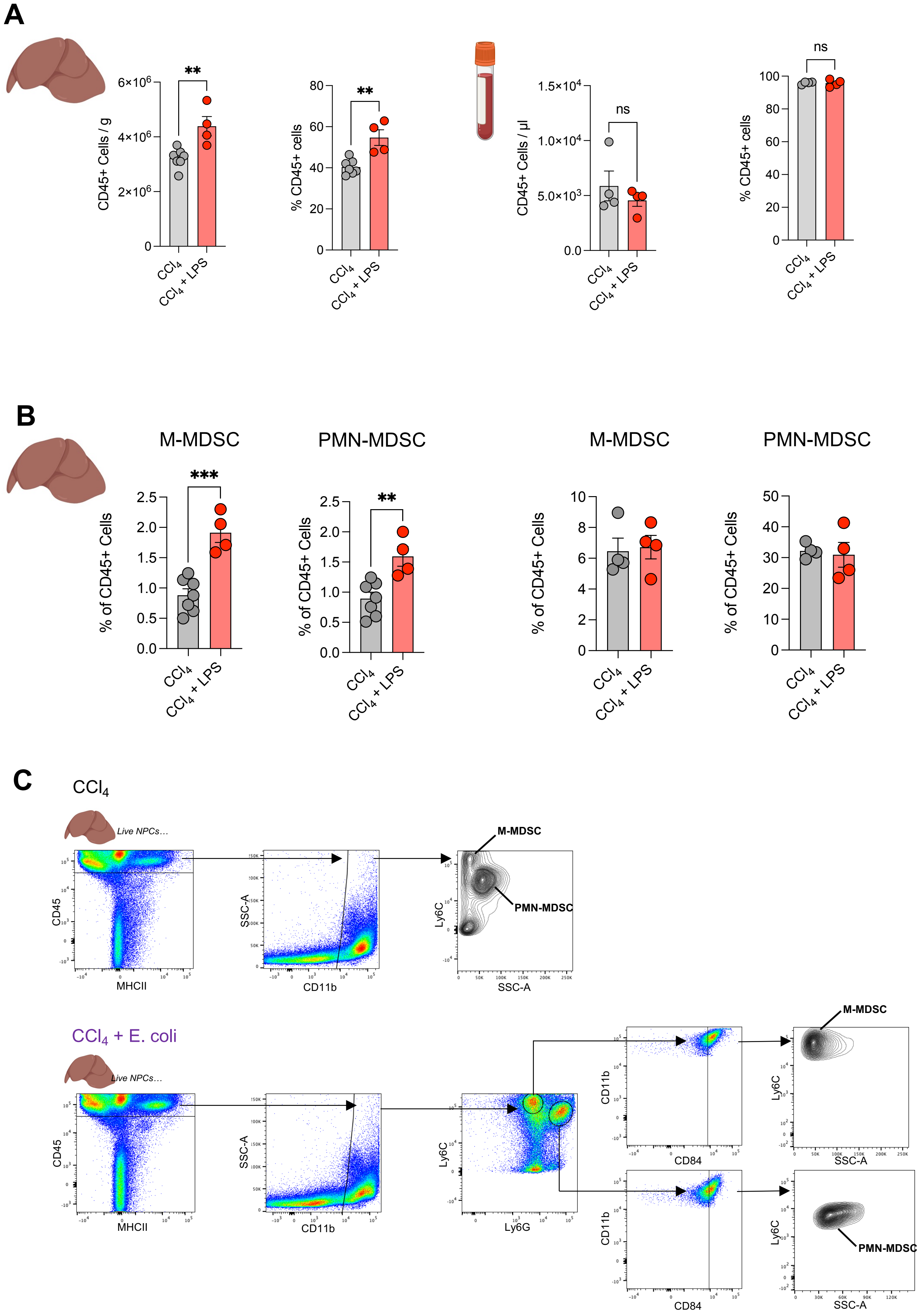

A
CD45+ Cells / g
6×10^6
4×10^6
2×10^6
0
**
CCl4
CCl4 + LPS
% CD45+ cells
80
60
40
20
0
**
CD45+ Cells / µl
1.5×10^4
1.0×10^4
5.0×10^3
0.0
ns
% CD45+ cells
100
80
60
40
20
0
ns
B
M-MDSC
PMN-MDSC
M-MDSC
PMN-MDSC
% of CD45+ Cells
***
**
2.5
2.0
1.5
1.0
0.5
0.0
10
8
6
4
2
0
50
40
30
20
10
0
CCl4
CCl4 + LPS
C
CCl4
Live NPCs…
CD45
MHCII
SSC-A
CD11b
Ly6C
SSC-A
M-MDSC
PMN-MDSC
CCl4 + E. coli
Live NPCs…
CD45
MHCII
SSC-A
CD11b
Ly6C
Ly6G
CD11b
CD84
Ly6C
SSC-A
M-MDSC
CD11b
CD84
Ly6C
SSC-A
PMN-MDSC

**Figure S3. (A)** Cell numbers and percentages of CD45+ cells (of total live cells) in both liver (left) and blood (right) compartments identified with flow cytometry analysis, cell numbers were normalised either to gram of liver tissue or to µl of blood using flow cytometry counting beads. Groups include $CCl_4$ controls and $CCl_4$-treated mice with LPS stimulation **(B)** Percentages of either M-MDSC or PMN-MDSC compared to total CD45+ cells in both liver (left) and blood (right) compartments identified with flow cytometry analysis. **(C)** Comparison of flow cytometry gating strategies, conventional (above) vs extensive (below) identification of MDSC subsets. Barplots displaying all datapoints with mean ± SEM, statistical tests by unpaired t tests, $^{**}p < 0.01$, $^{***}p < 0.001$.

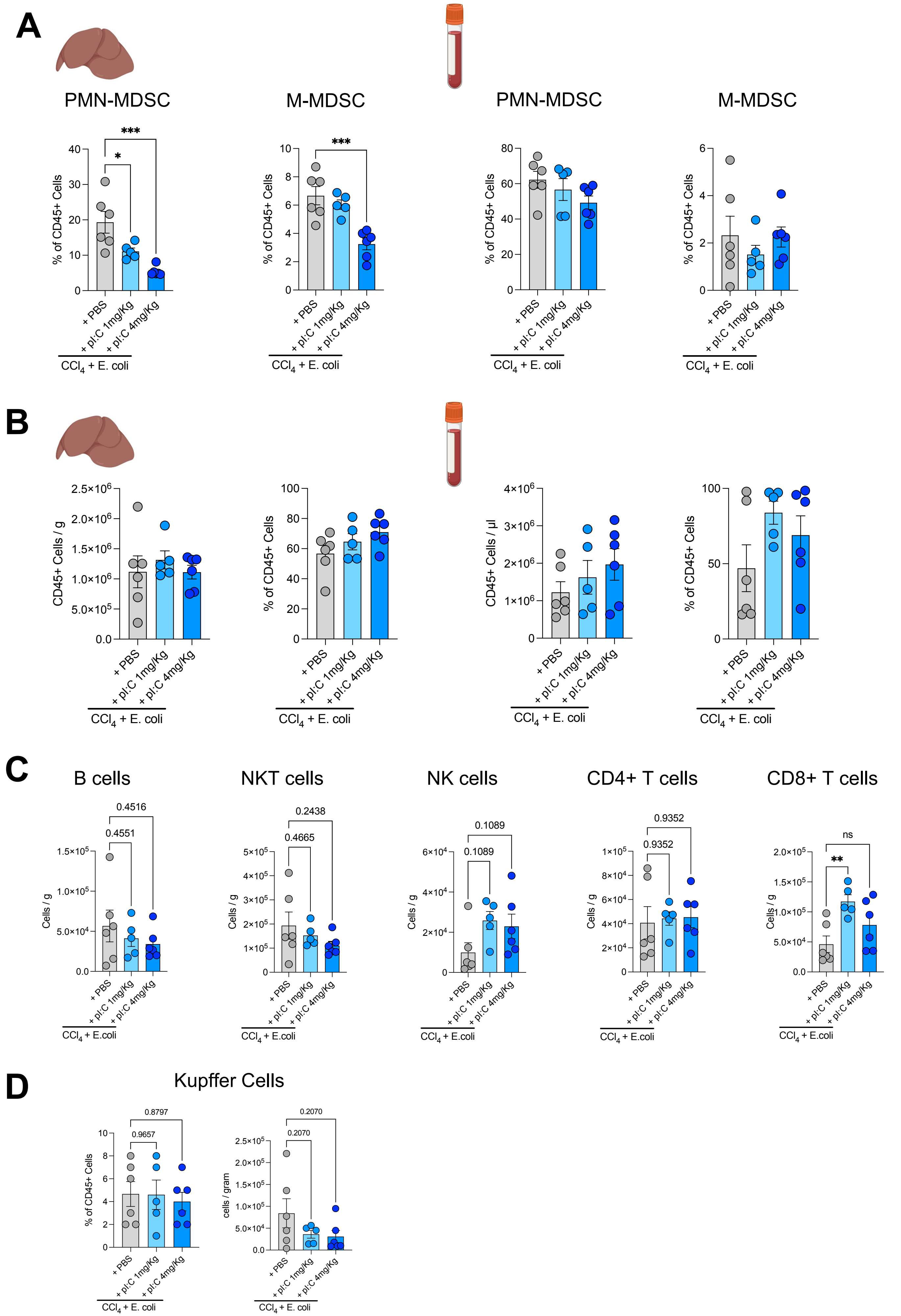
A
PMN-MDSC
M-MDSC
PMN-MDSC
M-MDSC
% of CD45+ Cells
*
***
+ PBS
+ pI:C 1mg/Kg
+ pI:C 4mg/Kg
CCl4 + E. coli
B
CD45+ Cells / g
% of CD45+ Cells
CD45+ Cells / µl
C
B cells
NKT cells
NK cells
CD4+ T cells
CD8+ T cells
Cells / g
0.4516
0.4551
0.2438
0.4665
0.1089
0.1089
0.9352
0.9352
ns
**
CCl4 + E.coli
D
Kupffer Cells
% of CD45+ Cells
cells / gram
0.8797
0.9657
0.2070
0.2070

**Figure S4. (A)** Percentages of either M-MDSC or PMN-MDSC compared to total CD45+ cells in both liver (left) and blood (right) compartments identified with flow cytometry analysis. Groups include $CCl_4$+infection controls, $CCl_4$+infection with poly(I:C) 1mg/Kg stimulation and $CCl_4$+infection with poly(I:C) 4mg/Kg stimulation. **(B)** Cell numbers and percentages of CD45+ cells (of total live cells) in both liver (left) and blood (right) compartments identified with flow cytometry analysis, cell numbers were normalised either to gram of liver tissue or to µl of blood using flow cytometry counting beads. **(C)** Cell numbers of hepatic lymphocyte subpopulations in $CCl_4$+infection controls, $CCl_4$+infection with poly(I:C) 1mg/Kg stimulation and $CCl_4$+infection with poly(I:C) 4mg/Kg stimulation. **(D)** Percentage of KC of total CD45+ immune cells and KC numbers per gram of liver tissue. Barplots displaying all datapoints with mean ± SEM, statistical tests by one-way ordinary ANOVA with multiple comparisons, ***$p < 0.001$.
